# Fast inactivation of Na^+^ current in rat adrenal chromaffin cells involves two independent inactivation pathways

**DOI:** 10.1101/2020.10.31.363341

**Authors:** Pedro L. Martinez-Espinosa, Alan Neely, Jiuping Ding, Christopher J. Lingle

**Author notes:** Department of Anesthesiology, Washington University School of Medicine, St. Louis MO USA 63105. Facultad de Ciencias, Centro Interdisciplinario de Neurociencia de Valparaiso, Instituto de Neurociencia, Universidad de Valparaıso, Valparaıso, Chile (Alan Neely). 203 Building 2, 17 Shatangyuan Road, Xuanwu, Nanjing, Jiangsu, China 210066 (Jiuping Ding). Correspondence to C. Lingle. **Competing interests**, The authors have no competing interests. **Author contributions**, P.L.M-E. A.N., and J.-P.D. contributed to whole-cell recording experiments, and designed and analyzed data. J.-P.D. was responsible for current-clamp experiments. C.J.L contributed to the design, analysis, interpretation of the results, and wrote the paper. All authors have approved the final version of the manuscript.

## Abstract

Voltage-dependent sodium (Nav) current in adrenal chromaffin cells (CCs) is rapidly inactivating and TTX-sensitive. The fractional availability of CC Nav current has been implicated in regulation of action potential (AP) frequency and the occurrence of slow-wave burst firing. To ascertain whether features of CC Nav inactivation might influence AP firing, we recorded Nav current in rat CCs, primarily from adrenal medullary slices. A key feature of CC Nav current is that recovery from inactivation, even following brief (5 ms) inactivation steps, exhibits two exponential components of generally similar amplitude. Variations of standard paired pulse protocols support the view that entry into the fast and slower recovery processes result from largely independent, competing inactivation pathways, both of which occur with similar onset times at depolarizing potentials. Over voltages from −120 to −80 mV, faster recovery varies from ~3 to 30 ms, while slower recovery from about 50-400 ms. At strong activation voltages (+0 mV and more positive), the relative entry into slow or fast recovery pathways is similar and independent of voltage. Trains of brief inactivating steps result in cumulative increases in the slower recovery fraction. This supports idea that brief recovery intervals preferentially allow recovery of channels from fast recovery pathways, thereby increasing the fraction of channels in the slow recovery pathway with each subsequent inactivation step. This provides a mechanism whereby differential rates of recovery produce use-dependent accumulation in slower recovery pathways. Consistent with use-dependent accumulation of channels in slow recovery pathways, repetitive AP clamp waveforms at 1-10 Hz frequencies reduce Nav availability to 10-20% of initial amplitude dependent on holding potential. The results indicate that there are two distinct pathways of fast inactivation, one that leads to normal fast recovery and the other with a slower time course, which together are well-suited to mediate use-dependent changes in Nav availability.

## Introduction

A classic view of the role of voltage-dependent Na^+^ (Nav) current is that it supports the reliable generation of action potentials (APs) of uniform duration and amplitude (Hille, 2001). This requires a sequence of rapid Nav current activation to produce cell depolarization, subsequent inactivation to help terminate net inward current, and then recovery from inactivation to permit a subsequent AP. The time course of recovery from rapid inactivation of Nav current contributes to a refractory period during which a cell is unable to generate a full AP (Hodgkin and Huxley, 1952; Kuo and Bean, 1994; Hille, 2001), potentially limiting cell firing rates. However, in many cells, recovery from fast inactivation is sufficiently rapid that repetitive AP firing can be sustained with little diminution in AP amplitude or change in AP frequency at AP frequencies exceeding 50 Hz (Schwindt et al., 1988; Wang et al., 1998; Khaliq et al., 2003; Kaczmarek et al., 2005; Brickley et al., 2007; Carter and Bean, 2011). However, in addition to fast inactivation, many Nav currents also exhibit an inactivation behavior in which recovery from inactivation occurs much more slowly, over hundreds of msecs or even seconds (Chiu, 1977; Rudy, 1981; Belluzzi and Sacchi, 1986; Jones, 1987; Ruff, 1996; Zhang et al., 2013; Silva, 2014). Such inactivation is sufficiently slow in onset that only in some unusual circumstances is it likely to influence Nav availability during normal firing (Silva, 2014).

However, over the past 15 years, the identification of additional Nav variants with distinct kinetic properties along with better elucidation of the complexity of Nav currents in native cells (Cummins et al., 1998; Dib-Hajj et al., 1999; Cummins et al., 2001; Hains et al., 2003; Herzog et al., 2003; Liu et al., 2003; Rush et al., 2006; Choi et al., 2007; Goldfarb et al., 2007; Milescu et al., 2010) has increased awareness that patterns of firing in some excitable cells may be influenced by use-dependent changes in availability of Nav channels. Furthermore, new mechanisms by which Nav channels can be regulated have been identified (Goldfarb, 2005; Rush et al., 2006; Goldfarb et al., 2007; Laezza et al., 2009; Shakkottai et al., 2009; Bosch et al., 2015). Specifically, for some Nav currents recovery from inactivation can occur at rates intermediate between traditional fast and slow recovery, involving a mechanism that appears distinct from either traditional fast or slow inactivation (Milescu et al., 2010; Goldfarb, 2012). This has been termed “long-term inactivation” (Dover et al., 2010; Barbosa and Cummins, 2016), which is distinguished from fast-inactivation by its relatively slower recovery from inactivation and is distinguished from slow inactivation by a rate of inactivation onset comparable to traditional fast inactivation. Long-term inactivation can be mediated by regulatory proteins termed intracellular fibroblast growth factor homologous factors (iFGFs) (Dover et al., 2010; Goldfarb, 2012; Venkatesan et al., 2014). Yet our understanding of such inactivation, the underlying relationship between different pathways of inactivation, and potential impact on cell firing remain rudimentary.

Here, we present results demonstrating that Nav current in rat adrenal chromaffin cells (CC) exhibits characteristics of so-called long-term inactivation. CCs offer a number of advantages over other cells for investigation of this phenomenon, including the simple spherical nature of the cells, the absence of processes, and a presumably a single type of rapidly inactivating, TTX-sensitive Na current (Lou et al., 2003; Vandael et al., 2015). The CC Nav current exhibits two distinct components of recovery from inactivation following rapid inactivation. In both mouse and rat CCs, Nav current availability has been proposed to impact on regulation of AP firing frequency (Solaro et al., 1995; Lingle et al., 1996), and slow wave burst firing (Vandael et al., 2015). Mouse and rat CCs fire APs in response to constant current injection at frequencies which rarely exceed 10-20 Hz (Solaro et al., 1995; Martinez-Espinosa et al., 2014). During such sustained depolarizing current injection, CCs typically exhibit a progressive decline in AP frequency or accommodation, in some cases leading eventually to block of AP firing. Although multiple factors (Lingle et al., 2018) may contribute to accommodation in CCs, including Ca^2+^-dependent SK-type K^+^ channels (Vandael et al., 2012) or properties of Ca^2+^-dependent BK-type K^+^ channels (Solaro et al., 1995; Martinez-Espinosa et al., 2014), use-dependent changes in Nav availability may also contribute. Careful definition of the properties of inactivation of rodent CC Nav current and its use-dependent alteration may therefore provide new insight into the role of Nav current in CC excitability.

Here we show Nav current in rat CCs exhibits two distinct components of recovery from fast inactivation, with both components entered at comparable rates during brief depolarizations. The two components differ by an order of magnitude in the rate of recovery from inactivation. There is little equilibration between pathways, once inactivation has occurred. Because rates of recovery differ between the two pathways, during repetitive stimuli channels exhibit use-dependent accumulation in the slower recovery pathway, which might contribute to slowing and eventual failure of AP generation. This dual-pathway, fast inactivation mechanism appears similar to that proposed to underlie inactivation mediated by the N-termini of intracellular fibroblast homologous growth factors (iFGFs) for Nav current in both hippocampal Purkinje cells (Venkatesan et al., 2014) and cerebellar granule cells (Goldfarb et al., 2007) and this is confirmed in the companion paper (Martinez-Espinosa et al., 2020). An analysis of Nav current in dorsal raphe neurons supports a similar dual-pathway model (Milescu et al., 2010), although the molecular basis for the inactivation in dorsal raphe neurons has not been determined. An accumulation of Nav channels in slow recovery states in raphe neurons has been proposed to act as a molecular integrator that essentially reports AP frequency and regulates firing (Navarro et al., 2020). A strength of the results presented here is that the rat CC currents are likely to arise from a single category of Nav channel, localized exclusively on the spherical plasma membrane. The use-dependent accumulation of Nav channels in the slower recovery pathway establishes a mechanism by which use-dependent changes in Nav availability can impact on cell firing behavior.

## Methods

### Animals

Sprague-Dawley rats were obtained from Harlan Laboratories or Jackson Labs. Rats were sacrificed by CO2 inhalation, following protocols approved by the Washington University in St. Louis Institutional Care and Use Committee. Following delivery of rats, animals were housed briefly in accordance with the National Institutes of Health Committee on Laboratory Animal Resources guidelines.

### Adrenal slice preparation

Adrenal glands from 8-12 week old rats were immediately removed following euthanasia and decapitation and immersed in ice-cold Ca^2+^- and Mg^2+^-free Locke’s buffer. Excess fat was trimmed from the glands, which were then embedded in 3% low gelling point agarose as described (Martinez-Espinosa et al., 2014). Agarose was prepared by melting agar in Locke’s buffer, followed by equilibration at 37°C. After embedding of tissue, the agarose block was trimmed to ~1-cm cubes, each containing a single gland, and then glued to a tissue stand of a vibratome (VT 1200 S; Leica). The tissue stand was then placed in a slicing chamber filled with ice-cold extracellular solution gassed with 95% O_2_/5% CO_2_. Glands were sectioned into 200-μm thick slices. Slices were collected and maintained in the gassed extracellular solution at room temperature until recording. All experiments were performed within 2–6 h after slice preparation.

### Slice recording methods and solutions

Recordings were done with a Multiclamp 700B (Molecular Devices, San Jose CA). The standard extracellular solution contained (mM) 119 NaCl, 23 NaHCO_3_, 1.25 NaH_2_PO_4_, 5.4 KCl, 2.0 MgSO_4_, 1.8 CaCl_2_, 11 glucose, 2 sodium pyruvate, and 0.5 ascorbic acid, pH 7.4. For most voltage-clamp recordings of Nav current, the traditional open pipette, whole-cell method was used with the pipette recording solution containing (in mM) 120 CsCl, 10 NaCl, 2 MgCl_2_, 5 EGTA, 1 Mg-ATP and 10 HEPES (pH 7.4 with CsOH). For open pipette experiments, 90% R_s_ compensation was typically employed.

Typical membrane capacitance for rat chromaffin cells included in this study was 9.0 ± 0.41pF. Pipette resistances ranged from 1.5 – 2.5 MΩ. Electrodes were coated with Sylgard 184 (Dow Chemical). Nominal voltages are uncorrected for R_s_ errors.

### Dissociated cell preparations and recording methods

Methods for isolation and long-term maintenance of CCs were as described (Neely and Lingle, 1992; Herrington et al., 1995) following methods developed by others (Kilpatrick et al., 1980; Role and Perlman, 1980; Livett, 1984). Such dissociated CCs were maintained in culture for 2-5 days.

Dissociated cells were used for perforated-patch recordings (Horn and Marty, 1988), in which spontaneous or evoked action potentials (APs) were recorded under current clamp and then the recorded AP waveforms were used as voltage-clamp commands. In perforated-patch recordings, for recording of APs, the pipette solution contained the following: 120 mM K-aspartate, 30 mM KCl, 10 mM HEPES(H+), 2 mM MgCl_2_ adjusted to pH 7.4 with N-methylglucamine (Herrington et al., 1995). For monitoring of AP-clamp evoked inward current, 20 mM KCl was replaced by equimolar CsCl in the pipette solution. Amphotericin B (Rae et al., 1991) and pluronic acid, stored in stock solutions of dimethylsulfoxide, were added to the pipette saline to make final concentrations of 500 μg/ml. The extracellular solution for such experiments was a standard physiological solution containing (in mM): 150 NaCl, 5.4 mM KCl, 2 mM CaCl2, 1 mM MgCl_2_, 5 mM HEPES, adjusted to pH 7.4. Recordings from dissociated cells were done with an Axoclamp 2A amplifier (Molecular Devices, San Jose CA).

### Solution exchange methods

Perfusion of external salines for dissociated cells was performed with a pipette with a single opening packed with six PE-10 internal perfusion lines, each under independent flow control (Herrington et al., 1995; Solaro et al., 1995). For adrenal slices, solution exchange was controlled from a single bore tube entering the slice chamber. All experiments were at room temperature (21-24°C).

### Data treatment and analysis

Voltage and current clamp commands and data acquisition was controlled with the Clampex program from the pClamp9 software package (Molecular Devices). Fitting of exponential relaxations and other functions was done with a non-linear least squares procedure utilizing the Levenberg-Marquardt algorithm or in some cases with function fitting procedures in Excel (Microsoft, Seattle USA). Statistical comparisons employed either a Student’s T-Test or a Kolgoromov-Smirnov (KS) test. Summary data are reported either as the mean ± STD of some number of individual cells (N) or, when averaged data are fit to a function, the value for the fitted parameter along with 90% confidence limit (c.l.) of that fitted parameter is provided.

## Results

### Basic properties of Nav current from CCs in rat adrenal medullary slices

Figure 1A shows a series of superimposed voltage-clamp records during 5 ms depolarizing steps of increasing amplitude following a 1000 ms step to −120 mV. Steps ranged from −70 mV to +50 mV in 5 mV increments, although only currents at 10 mV increments are displayed. Average peak current for 18 cells versus command potential is plotted in Figure 1B showing that I_Na_ peaks near −15 mV. For this set of cells, the average peak inward Nav current was −9494 + 2600 pA (mean±STD) corresponding to an average peak current density of −1067 pA/pF, which is several-fold larger than previously described for bovine (Fenwick et al., 1982) and mouse (Vandael et al., 2015) CCs. Assuming a +66 mV reversal potential, the normalized GV curves (Fig. 1C) for this set of cells yielded a voltage of half activation (Vh) of −27.4±0.2 mV with *z*=5.4±0.2*e*. This V_h_ is within about 5 mV of values reported for Nav current in rat sympathetic neurons (Belluzzi and Sacchi, 1986; Schofield and Ikeda, 1988) and mouse CCs (Vandael et al., 2015). The rate of onset of inactivation, measured from single exponential fits to the decay phase of the Nav current, was voltage-dependent with the time constant varying from about 2 ms at −25 mV, approaching 0.3 ms near 30 mV (Fig. 1D).

**Figure 1.**
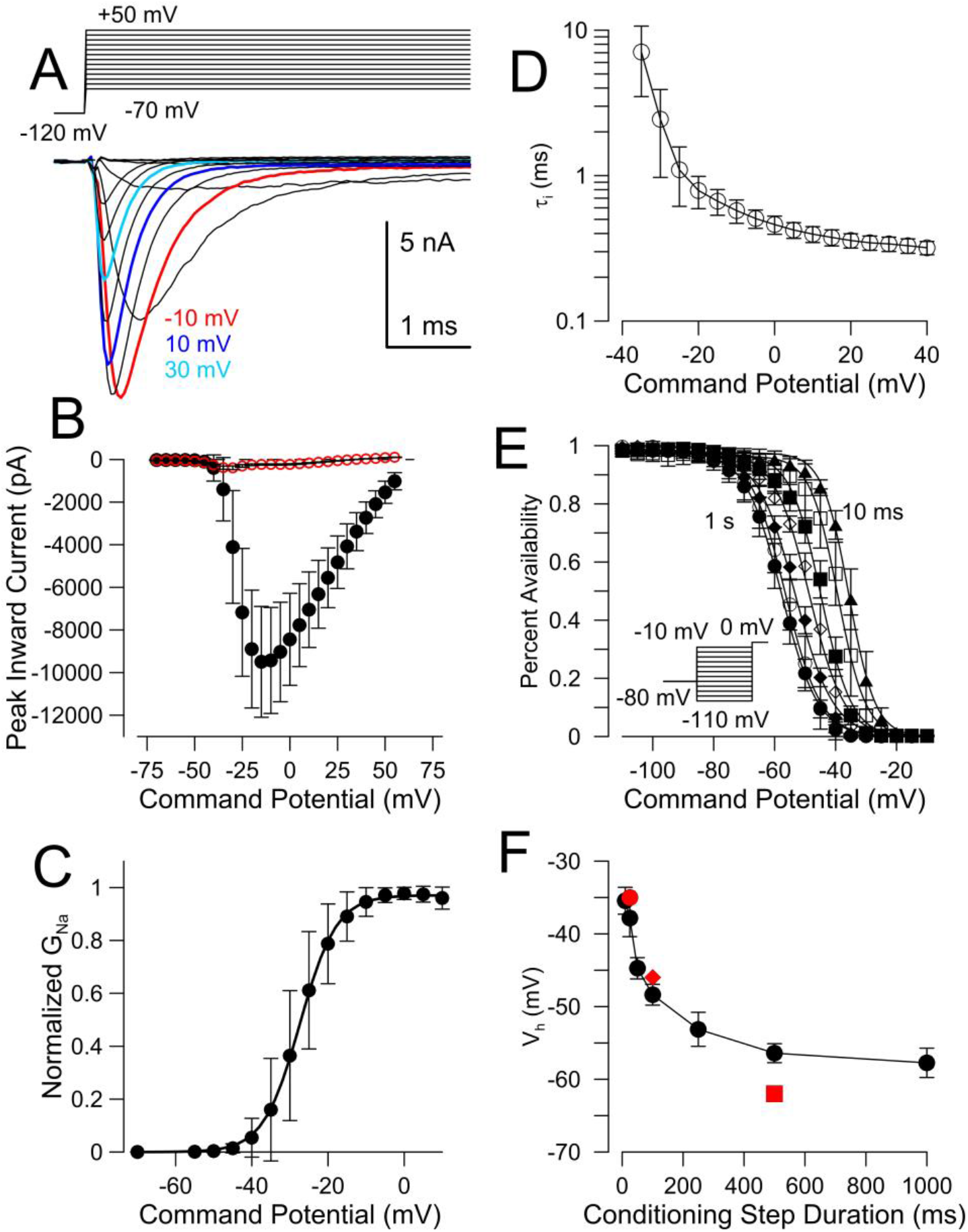
Basic properties of Nav current and steady-state inactivation in rat CCs. (A) Nav current was activated with the indicated stimulus protocol in a rat adrenal CC in a slice. (B) The averaged peak-current density for inward current is plotted (mean± STD) for a set of 18 rat CCs. The steady-state current (at end of 5 ms step) for the same set of cells is shown in red. (C) Currents from the set of patches in panel B were used to generate a G/V curve (means±STD), assuming ENa= 66 mV. For each cell, extrapolation of currents activated at +50 and +55 mV yielded ENa=64.4±2.9 mV (mean±STD), while ENa calculated from nominal intracellular and extracellular solutions was 67 mV. The fitted voltage of half activation was −27.4 ± 0.2 mV with z=5.4±0.2*e*. (D) The rates (means±STD; n=12 cells) of onset of inactivation measured from single exponential fits to the decay phase of the Nav currents are plotted as a function of the inactivation voltages. (E) Fractional availability (means±STD) is plotted as a function of conditioning potential over a range of conditioning potential durations (10, 25, 50, 100, 250, 500, 1000 ms). Inset shows example voltage protocol, where conditioning potential ranged from −110 to −10 mV (5 mV increments), with a test potential of 0 mV, following an initial holding potential of −80 mV. Number of patches tested for each recovery voltage were, from 10 to 1000 ms, 8, 4, 6, 9, 6, 7, and 13. V_h_ and *z* for single Boltzman fits were, for 10 ms, −35.6±0.3 mV, *z*=5.9±0.3*e*; for 25 ms, V_h_=-39.2±0.4 mV, *z*=5.7±0.4*e*; for 50 ms, V_h_=-44.8±0.4 mV, *z*=5.1±0.4*e*; for 100 ms, V_h_=-48.7±0.5 mV, *z*=4.5±0.3*e*; for 250 ms, V_h_=-53.2±0.4 mV, *z*=4.1±0.2*e*; for 500 ms, V_h_=-56.5±0.3 mV, z=4.4±0.2; for 1000 ms, V_h_=-57.7±0.2 mV, *z*=4.3±0.1*e*. (F) The mean V_h_ of fractional availability obtained from single Boltzman fits to the curves from each individual cell used for the averages in panel E is plotted (±STD; n=4-13 cells, as indicated) as a function of conditioning potential duration. The line simply connects the dots. Red symbols correspond to measurements of voltage of half availability in previous papers (see text).

In bovine CCs, the voltage at which Nav channels are half inactivated at steady-state was reported to be −34 mV using 25 msec conditioning steps (Fenwick et al., 1982). In contrast, one study of rat CCs, using a 250 ms conditioning step, reported a V_h_ for steady-state inactivation of –60 mV (Hollins and Ikeda, 1996), while another reported a V_h_ of −62 mV with a 500 ms prepulse duration (Lou et al., 2003). Here, we examined the effect of conditioning duration on the V_h_ of inactivation. With a 25 msec conditioning step, half-inactivation was observed at −35.4 mV (Fig. 1E-F), similar to that reported for bovine CCs (Fenwick et al., 1982). However, the steady-state inactivation curves shifted considerably to more negative voltages as the duration of the conditioning step duration increased from 25 msec up to 1 sec (Fig. 1E), with a V_h_ for steady-state inactivation of −57.7 mV with the 1 sec conditioning step (Fig. 1F). The modest change in V_h_ between 500 ms and 1 sec suggests that a conditioning step of at least 1 sec is necessary to approach the equilibrium voltage-dependence of Nav availability in rat CCs.

### Recovery from inactivation, even following brief inactivation steps, occurs with both fast and slow components

The temporal features of recovery from inactivation were examined using standard paired depolarizing pulses to 0 mV (Fig. 2A) separated by recovery periods of varying length at a given recovery potential. An initial holding potential of −80 mV, at which more than 95% of the Nav channels are available for activation (Fig 1E), preceded the first test pulse (P1). I_Na_ amplitude during the second depolarizing pulse (P2) following each recovery interval was normalized to I_Na_ activated during P1. Example currents evoked from the paired pulse stimulation protocol (1 ms to 3 s recovery) are shown in Figure 2A,B with recovery either at −60 mV (Fig. 2A) or −120 mV (Fig. 2B). Over a range of recovery potentials (−60 to −120 mV), the fractional recovery as a function of recovery duration is not well-described by a single exponential time course (Fig. 2C), while a double exponential provides an excellent fit (Fig. 2D). The double exponential nature of the recovery time course can be best seen by plotting the recovery durations on a logarithmic time scale (right-hand panels of Fig. 2C,D).

**Figure 2.**
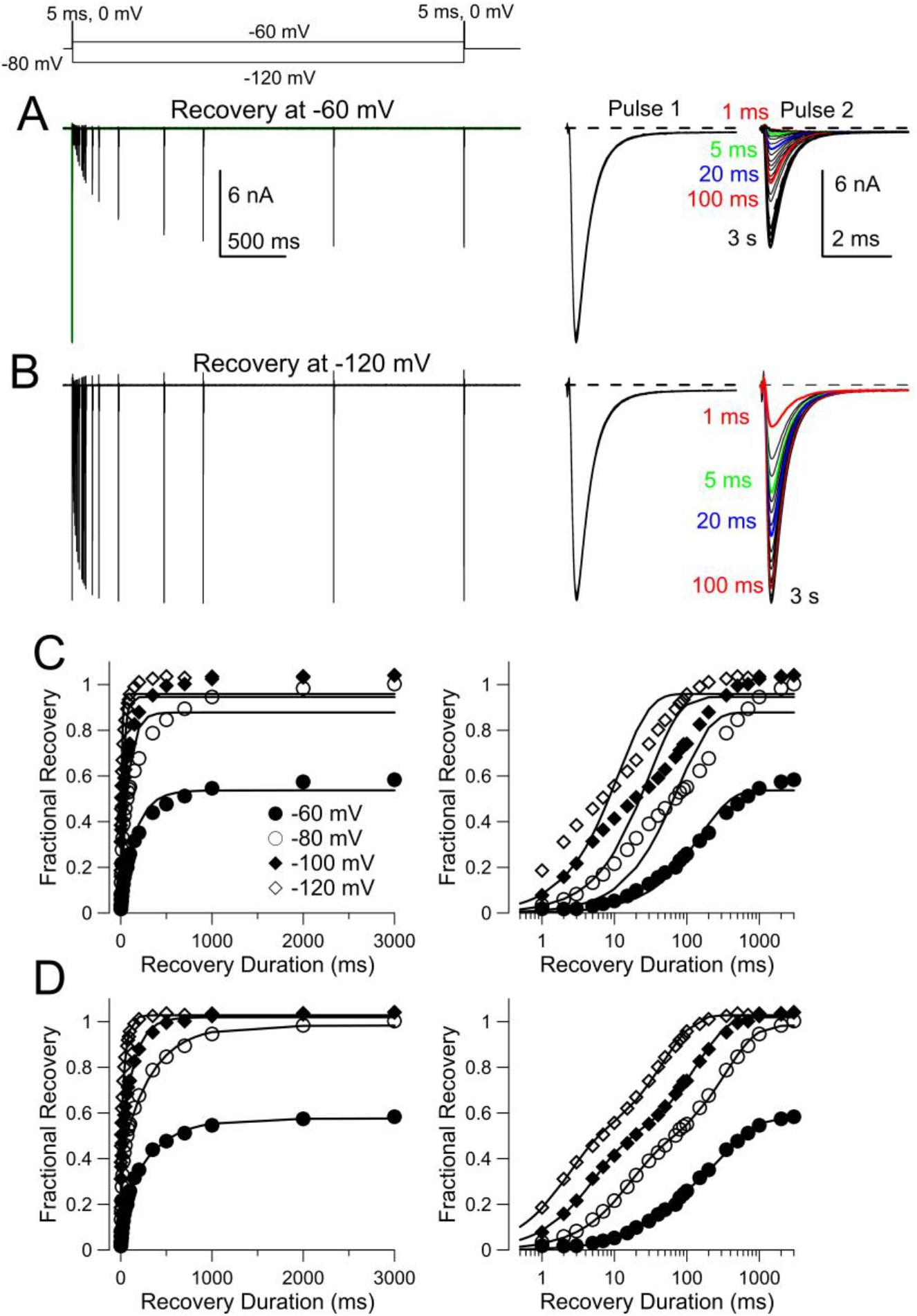
Recovery from fast inactivation involves two components. A paired pulse protocol (shown on top) was used to examine the time course of recovery from inactivation. From a holding potential of −80 mV, a 5 ms activation step to 0 mV was used to produce Nav inactivation. In a given trial, the cell was then repolarized to a voltage between −60 and −120 mV for durations from 1 ms to 3 s before the second test step to 0 mV. The example protocol (top) shows cases for 3 s recovery at either −60 or −120 mV. (A) A family of traces showing recovery from inactivation at −60 mV. On the right, traces are shown for the response to the initial step to 0 mV (pulse 1: P1) and then for the response to the second step to 0 mV (pulse 2: P2) following the recovery step. Colored traces highlight recovery at the indicated intervals. (B) As in A, traces are shown for recovery at −120 mV. (C) Fractional recovery from 1 ms to 3 s is plotted for recovery at −60, −80, −100, and −120 mV for the cell shown in A and B using a linear scale on the left and a logarithmic scale on the right. Lines are fits of a single exponential function to the recovery time course: *I_t_* = *A*(1−exp(−*t*/τ)) where *A* and τ are the amplitude and time constant of the recovery process, respectively. (D) The same fractional recoveries shown in C are replotted along with fits of a double exponential function to the recovery time course with *I_t_* = *A_f_* (1−exp(−*t*/τ_f_)) ± *A_s_* (1− exp(−*t* /τ_s_)). For −60 mV, *A*_f_=0.19±0.03, *τ_f_*=49.9±7.9 ms, *A_s_*=0.38±0.02, τ_s_=373.7±34.6 ms (fitted value and 90% confidence limit). For −80 mV, *A*_f_=0.40±0.01, *τ_f_*=14.8±0.9 ms, *A_s_*=0.59±0.01, τ_s_=332.0±19.7 ms. For −100 mV, *A_f_*=0.44±0.01, τ_*f*_=5.0±0.4 ms, *A_s_*=0.58±0.1, τ_s_=134.7±8.4 ms. For −120 mV, *A_f_*=0.46±0.01, *τ_f_*=2.1±0.1 ms, *A_s_*=0.57±0.01, τ_s_=46.1±2.4 ms.

The averaged fractional recovery and associated standard errors were plotted along with the best fit of the double exponential function for one set of cells studied at recovery potentials of −60, −80, −100, and −120 mV (Fig. 3A) and another set of cells at −50, −70, −90, and −110 mV (Fig. 3B). Over the tested recovery voltages, the faster time constant of recovery ranged from about 55.9 ± 1.1 ms at −60 mV to 2.9 ± 0.4 ms at −120 mV, while the slower time constant plateaued at 487 ± 8 ms at −60 mV, and was around 56.5 ± 0.6 ms at −120 mV (Fig. 3C). A plot of the relative amplitude of each recovery component as a function of voltage shows that, at potentials negative to about −80 mV, most inactivated channels fully recover from inactivation (Fig. 3D). However, at more positive recovery potentials, the relative amplitudes of the recovery components through slow and fast recovery components exhibit some differences. Thus, with recovery at −70 mV, the absolute amplitude of the slow recovery process is similar to that at the most negative recovery potentials, but the amplitude of the fast recovery component is reduced.

**Figure 3.**
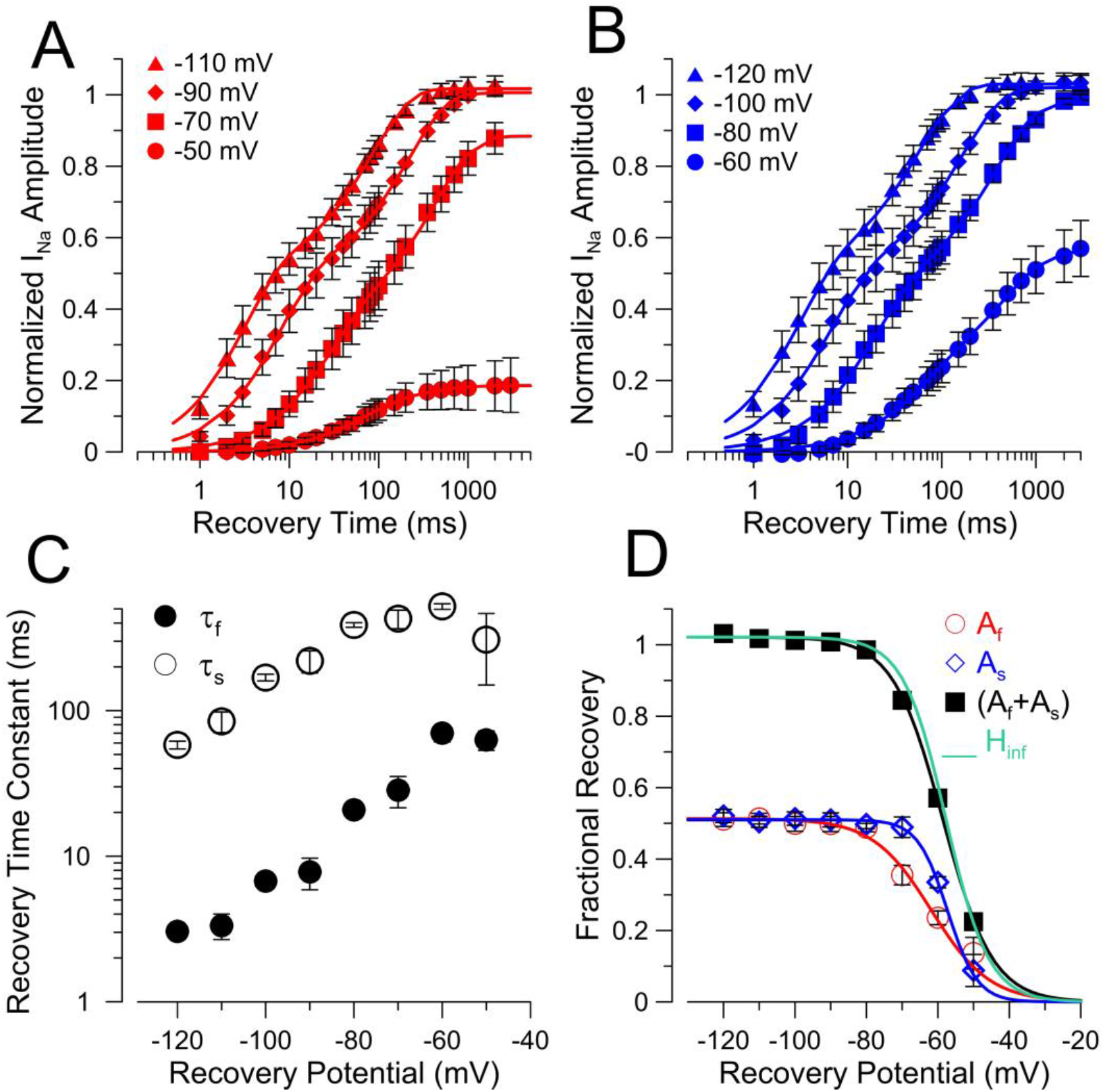
Voltage-dependence of two components of recovery from inactivation. (A) A log-scale plot of averaged fractional recovery from inactivation following a 5 ms depolarizing voltage step to 0 mV is shown for recovery voltages of −50, −70, −90 and −110 mV, using the protocol illustrated in Fig. 2. Each point shows the mean ± STD (at −50 mV, n=11 cells; at −70, n=14; at −90, n=15; at −110 mV, n=13. Lines are the best fit of a two exponential function to the recovery time course. Each cell was equilibrated at −80 mV for 1 s prior to the initial inactivating test step to 0 mV. At −110 mV, *A_f_*=0.51±0.01, *τ_f_*=3.1±0.2 ms, *A_s_*=0.51±0.01, τ_s_=83.2±4.7 ms (fitted value and 90% confidence limit); at −90 mV, *A_f_*=0.50±0.01, *τ_f_*=7.6±0.5 ms, *A_s_*=0.51±0.01, τ_s_=215.3±17.1 ms; at −70 mV, *A_f_*=0.36±0.01, *τ_f_*=23.1±1.6 ms, *A_s_*=0.53±0.13, τ_s_=404.9±27.3 ms; for −50 mV, *A_f_*=0.14±0.01, *τ_f_*=64.7±5.8 ms, *A_s_*=0.05±0.01, τ_s_=359.1±110.9 ms. (B) For a different set of cells (10 or 11 cells), fractional recovery was determined at −60, −80, −100, and − 120 mV, with solid lines indicating the double exponential fit. At −120 mV, *A_f_*=0.51±0.01, *τ_f_*=2.9±0.4 ms, *A_s_*=0.52±0.01, τ_s_=56.5±0.6 ms; at −100 mV, *A_f_*=0.50±0.01, *τ_f_*=6.4±0.6 ms, *A_s_*=0.53±0.01, τ_s_=162.9±1.8 ms; at −80 mV, *A_f_*=0.45±0.01, *τ_f_*=18.4±0.3 ms, *A_s_*=0.53±0.01, τ_s_=373.5±6.1 ms; at −60 mV, *A_f_*=0.21±0.01, *τ_f_*=55.93±1.1 ms, *A_s_*=0.34±0.01, τ_s_=487.8±7.6 ms. (C) At each recovery voltage, the mean fast and slow time constants (±STD) obtained from fits to individual cells are plotted for each set of cells. These mean values for sets of individual cells agree closely with the fits to the averaged data shown in A and B. (D) The mean amplitude (±STD) of faster and slower recovery components are plotted as a function of voltage along with the sum of the two components. Single Boltzmann fits to each relationship are shown. For the fast component (red circles), Af = 0.51±0.02, V_h_=-61.8±1.5 mVand z=3.2±0.8*e*; for the slow component (blue circles), A_s_=0.51±0.01, V_h_=-57.3±0.87 mV, and *z*=6.2±1.9*e*. For the sum of the two components (black rectangles), V_h_=-58.6±0.3 mV with *z*=3.7±0.2*e*, agreeing closely with the measurements of fractional availability in Fig. 1E (green line), V_h_=-57.7 mV and *z*=4.3*e*.

Minimally, the results indicate that multiple inactivated states participate in the recovery process, but interpretation of the significance of the slow and fast recovery components depends on the relationship between those inactivated states. For the moment, we propose that the two components reflect largely separate and independent inactivation pathways and we test this further below.

Empirically, the fraction of channels recovering either through slow or fast pathways as a function of recovery potential can each be approximated by a Boltzmann function 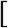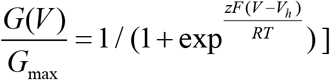. For the fast recovery component, V_h_ is −61.8 ± 1.5 mV (*z* =3.1±0.8*e*), whereas, for the slow recovery component, the V_h_ is −57.3 ± 0.9 mV (z= 6.2±1.9*e*). When the amplitude of both recovery components are added together, reflecting the full fractional recovery at each recovery potential, the summed amplitudes can be well-fit with a single Boltzmann with V_h_ = −58.6±0.3 mV with *z*=3.7±0.2*e*, agreeing closely with the steady-state inactivation curves measured following 1 sec conditioning potential (Fig. 1E).

Although we have not examined this in detail, we did observe some cells for which the relative amplitudes of the fast and slow components of recovery were not as similar as for the cells in Fig. 3, e.g., Figs. 4-5. We have no definitive answer for such differences in A_f_ and A_s_. Possible explanations might include: 1. differential expression of two populations of Nav channels; 2. differential contribution of some molecular component that contributes to the slower component of recovery from inactivation. We consider the latter possibility more likely (Martinez-Espinosa et al., 2020).

**Figure 4.**
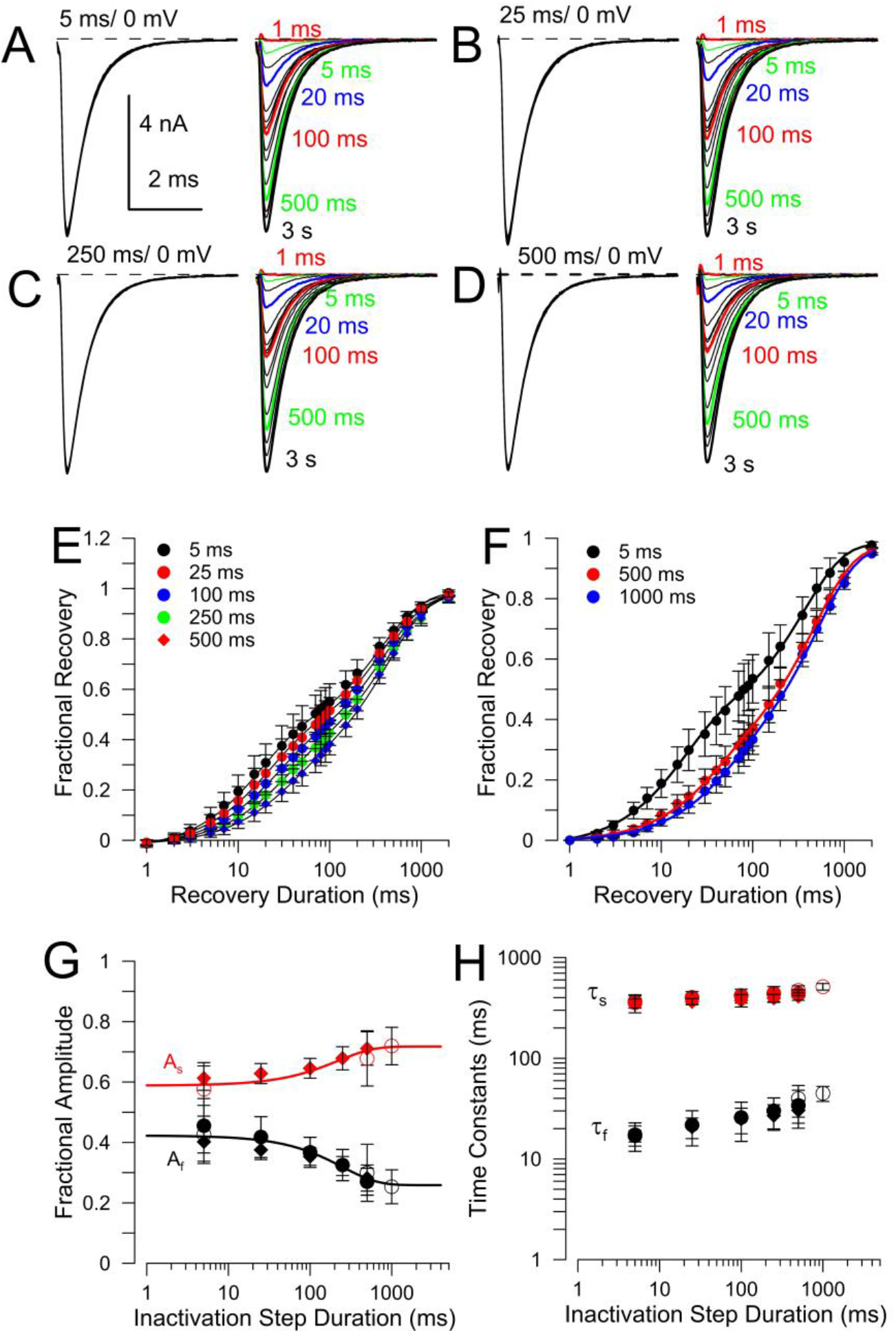
Effects of variation in duration of inactivation step on relative amplitudes of fast and slow components. (A-D) Paired depolarizing pulses to ±0 mV from a −80 mV holding potential were used to examine the time course of recovery from inactivation with inactivation pulse durations of 4, 25, 250, and 500 ms duration. All traces are from the same cell, with colored traces showing recovery durations preceding the 2^nd^ pulse indicated on each panel. (E) Mean fractional recovery (16 cells for 5, 25, 100, and 250 ms, and 10 cells for 500 ms) following the indicated inactivation durations are plotted as a function of recovery duration, with each set of points fit by a double exponential recovery function. Errors bars in E and F are STD. For 5 ms inactivation, *A_f_*=0.44±0.01, *τ*_f_=16.2±1.12 ms, *A_s_*=0.57±0.01, τ_s_=359.4±21.3 ms; for 25 ms, *A_f_*=0.41±0.01, *τ_f_*=19.4±1.5 ms, *A_s_*=0.60±0.01, τ_s_=389.9±22.2 ms; for 100 ms, *A_f_*=0.36±0.01, *τ_f_*=22.7±1.8 ms, *A_s_*=0.63±0.01, τ_s_=404.6±20.3 ms; for 250 ms, *A_f_*=0.32±0.02, *τ*_f_=27.9±3.1 ms, *A_s_*=0.68±0.02, τ_s_=430.3±24.7 ms; for 500 ms, *A_f_*=0.26±0.01, *τ_f_*=31.1±3.5 ms, *A_s_*=0.73±0.01, τ_s_=432.3±18.9 ms. (F) Recovery was examined in another set of 4 cells for 5, 500, and 1000 ms inactivation durations at 0 mV, again with double exponential fits overlaid. For 5 ms, *A_f_*=0.40±0.01, *τ_f_*=16.4±1.4 ms, *A_s_*=0.60±0.01, τ_s_=364.9±22.6 ms; for 500 ms, *A_f_*=0.26±0.02, *τ_f_*=34.6±3.4 ms, *A_s_*=0.71±0.01, τ_s_=476.1±23.1 ms; for 1000 ms, *A_f_*=0.24±0.02, *τ_f_*=44.5±5.9 ms, *A_s_*=0.73±0.02, τ_s_=509.2±28.9 ms. (G) The amplitude of the fast (Af) and slow (As) recovery components are plotted as a function of recovery duration. In this case, the Af and As values from each individual cell in panel E were averaged to determine the mean ± STD Filled circles correspond to cells tested sequentially with 5, 25, 100, 250, and 500 ms inactivation durations. Filled diamonds correspond to cells tested in the sequence of 500, 250, 100, 25, and 5 ms. Open circles correspond to four cells tested with 5, 500, and 1000 ms durations in both forward and reverse orders. Solid lines correspond to single exponential fits to the temporal changes in each amplitude component. Changes in Af occur with a time constant of 248.0 ± 126.0 ms, while As changes with a time constant of 219.0 ± 205.1 ms. Af, As, *τ_f_*, and τ_s_ values measured at 5 and 25 ms were not statistically different. (H) The time constants for fast and slow recovery monitored at −80 mV following inactivation steps for different durations to 0 mV are plotted, illustrating a slow prolongation of both *A_f_* and τ_s_ with inactivation duration. Errors bars are STD.

### Models of slow recovery from inactivation

To guide consideration of the experiments presented below, we begin with two general categories of scheme by which multiple components of recovery from inactivation might occur. Scheme 1 encapsulates a traditional and general view of slow inactivation (Zhang et al., 2013; Silva, 2014) in which there are two tiers of inactivated states, one (I_1_-I_6_) in which recovery from inactivation is fast, and another (I_1_’-I_6_’) in which return to other states is slow. Thus, occupancy of more slow recovering states occurs sequentially from occupation of fast recovering states. More recent work suggests that movement of the Domain IV voltage-sensor of Nav1.4 is rate limiting for fast inactivation(Capes et al., 2013), but for present purposes Scheme 1 encapsulates a general category of model in which traditional slow inactivation is linked to entry into fast inactivated states.

**Scheme 1.**
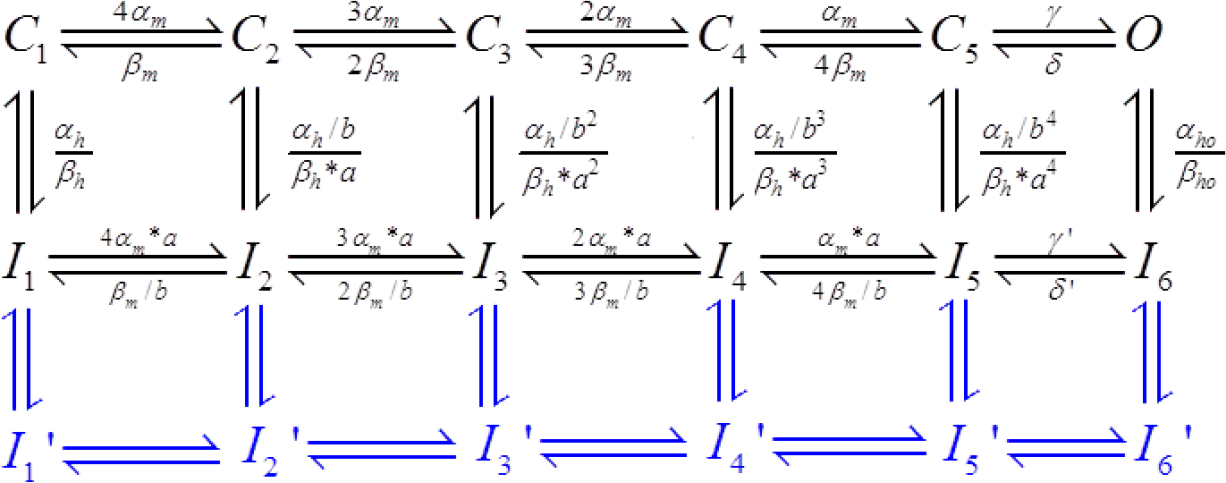

Scheme 1 requires that entry into states from which recovery occurs slowly depends first on entry into the fast recovery states. A characteristic of such a scheme is that, as the duration of the inactivation step is increased, the amplitude of any slow component of recovery will increase (Zhang et al., 2013; Silva, 2014). It should be noted immediately that the observation in CCs that both slow and fast components of recovery of about equal amplitude are observed following a 5 ms depolarization is inconsistent with Scheme 1, unless entry into I_1_’-I_6_’ is very rapid.

A second general model has been proposed to explain the effects of intracellular fibroblast growth factor homologous factors (iFGFs) on slow recovery from inactivation of particular Nav currents in neurons (Goldfarb et al., 2007; Dover et al., 2010; Milescu et al., 2010; Goldfarb, 2012; Venkatesan et al., 2014). This scheme (Scheme 2) posits two distinct fast inactivation pathways, one identical to normal fast inactivation (I_1_-I_6_) while the other, I_SR_, involves a competing fast inactivation pathway into states that recover more slowly (SR) from inactivation. Scheme 2 also includes traditional slow inactivation states. As schematized in Scheme 2, the secondary fast inactivation pathway only occurs from the open state, but this is only for illustrative purposes and this category of mechanism might also permit inactivation from some number of closed states (Milescu et al., 2010; Venkatesan et al., 2014). An important idea encapsulated in Scheme 2 that distinguishes it from Scheme 1 is that entry into ISR is essentially competitive with normal fast inactivation, under conditions that favor channel opening. Conceivably, other mechanisms might allow the possibility of transitions between the two distinct inactivation pathways, e.g., between I_SR_ and I_6_, but as shown below this does not seem required by our data.

**Scheme 2.**
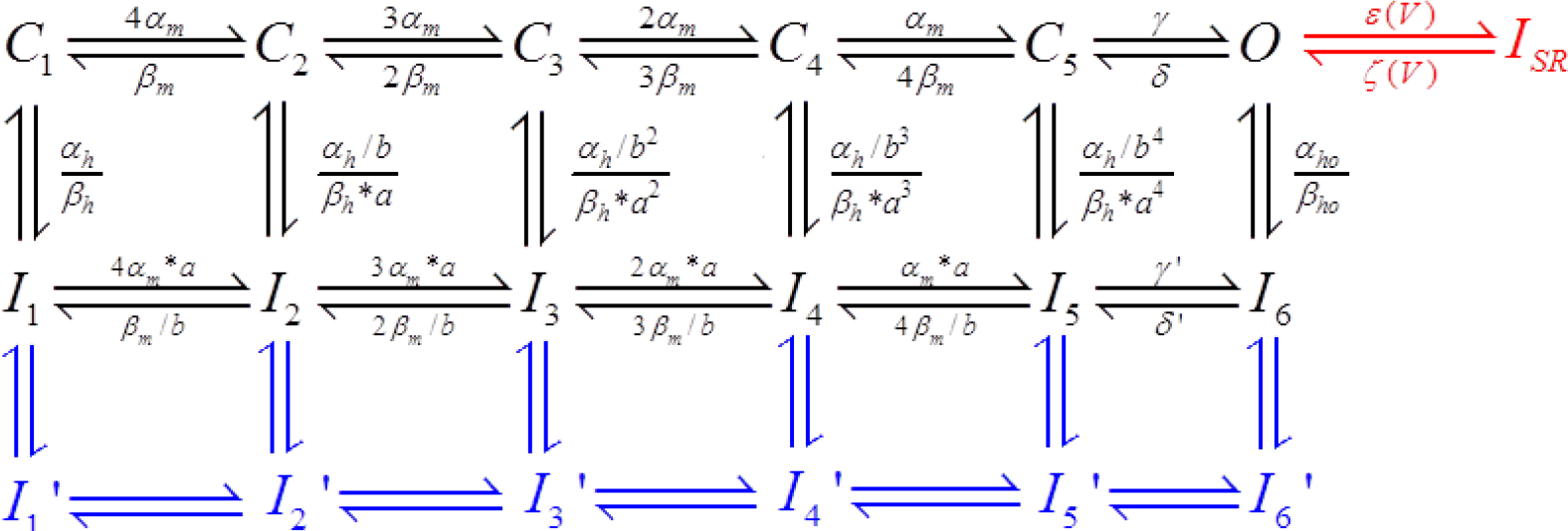

Although both Schemes 1 and 2 are consistent with currents that exhibit both slow and fast components of recovery from inactivation, qualitatively Schemes 1 and 2 make very different predictions for the properties of entry and recovery from inactivation. We now utilize a set of protocols to show that inactivation of Nav current in rat CCs exhibits a behavior consistent with Scheme 2, but not Scheme 1. The basic approach is to utilize protocols that, beginning from some specific initial condition, e.g., channels in resting states or channels in some condition of steady-state inactivation, we then monitor either the time course of recovery from inactivation or onset of inactivation, in order to try to tease apart the connectedness between states underlying the two observed inactivation time constants.

### The duration of an inactivation step does not strongly influence distribution between slow and fast recovery paths at strong inactivation voltages

For traditional slow inactivation, as the duration of a strong inactivating depolarization is increased, the relative amplitude of slow recovery from inactivation gradually increases (Zhang et al., 2013; Silva, 2014). At first glance, the fact that both fast and slower recovery from inactivation occurs with comparable fractional amplitudes after even a 5 ms inactivation step tends to argue against Scheme 1, i.e., traditional slow inactivation. To examine this more closely, we determined the relative amplitude and rates of fast and slow recovery from inactivation using paired pulses to +0 mV, but with inactivation durations of 5, 25, 100, 250, and 500 ms (Fig. 4A-D; Fig. 4E). The recovery time courses from one cell show that, as the duration of the inactivation step is increased, there is some decrement in the amplitude of the fast recovery component at longer inactivation durations (compare Fig. 4A to Fig4C-D). Plotting the fractional recoveries for a set of 16 cells tested at 5, 25, 100, and 250 ms for which 10 of the cells were also tested at 500 ms showed a consistent small decrement in the fast component (Fig. 4E) and some slowing of both τ_f_ and τ_s_. Because of concern that the sequence of the pulse protocols might result in some slow change in channel behavior, we also used a reverse sequence of inactivation durations of 500, 250, 100, 25, and 5 ms in a set of 4 cells and obtained similar changes in the amplitudes and durations of recovery components with longer inactivation durations (plotted in Fig. 4G). Longer inactivation durations of (500 and 1000 ms) were explored in another set of 4 cells (Fig. 4F). The changes in amplitudes of fast (A_f_) and slow (A_s_) recovery components were plotted as a function of the inactivation step duration (Fig. 4G) and fit with a single exponential function, indicating that the decrement in A_f_ occurred with a time constant of 248 ± 126 ms approaching steady-state at a fractional amplitude of 0.26 ± 0.3, while the increase in A_s_ occurred with a similar time constant of 219 ± 205 ms and limiting steady-state amplitude of 0.72 ± 0.02. Both the fast and slow recovery time constants were slightly slowed with longer inactivation step durations (Fig. 4H). Whatever the origin of the changes in the properties of the fast and slow recovery components, they occur with time constants much slower than the initial rapid rates of entry into the two recovery pathways. The overall result indicates that, once channels have distributed about equally between the two recovery pathways after a 5 ms depolarization, there is only a small additional change in relative fractions of fast and slower recovering channels up through 100 ms at the inactivating voltage.

What might be the origin of the small change in ratio of fast and slow components of recovery with prolonged inactivating pulses? Although it is possible there is some additional equilibration between the two pathways after the initial fast inactivation, another possibility is that it might simply reflect a small contribution of something akin to traditional slow inactivation (Zhang et al., 2013; Silva, 2014). For Nav1.5, a TTX-resistant cardiac Nav channel for which some quantitative estimates of slow inactivation are available (Zhang et al., 2013), entry into slow inactivation at −20 mV was observed to occur with a time constant of 1.8 sec, with slow inactivation only beginning to be detectable with inactivation durations of 100 or 200 ms. Recovery from slow inactivation for Nav1.5 occurs with time constants on the order of hundreds of ms at −120 mV. The onset of Nav1.5 slow inactivation is slower than what we observe here for the additional slow equilibration of the CC Nav currents. However, overall the slow changes in the amplitudes of both the slow and fast components and the prolongations in the time constants are generally consistent with what would be expected, if channels were entering an additional slow recovery state. However, the present results do not provide any definitive insight into this issue.

Although it might be tempting to infer that the two recovery time constants for inactivation durations up to 100 ms reflect distinct inactivation pathways for which there is little equilibration following inactivation (Scheme 2), Scheme 1 could apply under the condition that equilibration betweem the two tiers of inactivated states in Scheme 1 has to be complete within 5 ms and not change substantially change over 100 ms. If so, that would be a highly unusual form of traditional slow inactivation.

### Relative entry into slow and fast recovery paths exhibits little voltage-dependence above 0 mV

We next examined whether relative entry into the two pathways might differ at different inactivation voltages. We focused on voltages over which activation of Nav current was near saturation (Fig. 1C) such that any differences would largely might reflect intrinsic voltage-dependence in transitions involved in inactivation. Recovery at −80 mV was examined following inactivation produced by a 25 ms step at command voltages from −10 to +30 mV (Fig. 5A). An inactivation step of 25 ms was chosen since inactivation is essentially complete within this time period over these voltages. As seen during the currents activated during P2, at inactivation voltages from −10 to +30 mV, there is little change in the time course of recovery from inactivation (Fig. 5A-B). After a recovery duration of 100 ms, a similar fraction of channels have recovered from inactivation irrespective of the inactivation potential. For a set of over 10 cells, the amplitude of the fast component was slightly diminished at −10 mV but essentially constant from 0-30 mV (Fig. 5C-D). The inactivation voltages over which the relative amplitudes of the two recovery components are voltage-independent corresponds approximately to voltages over which Nav conductance is maximally activated (Fig. 1C).

**Figure 5.**
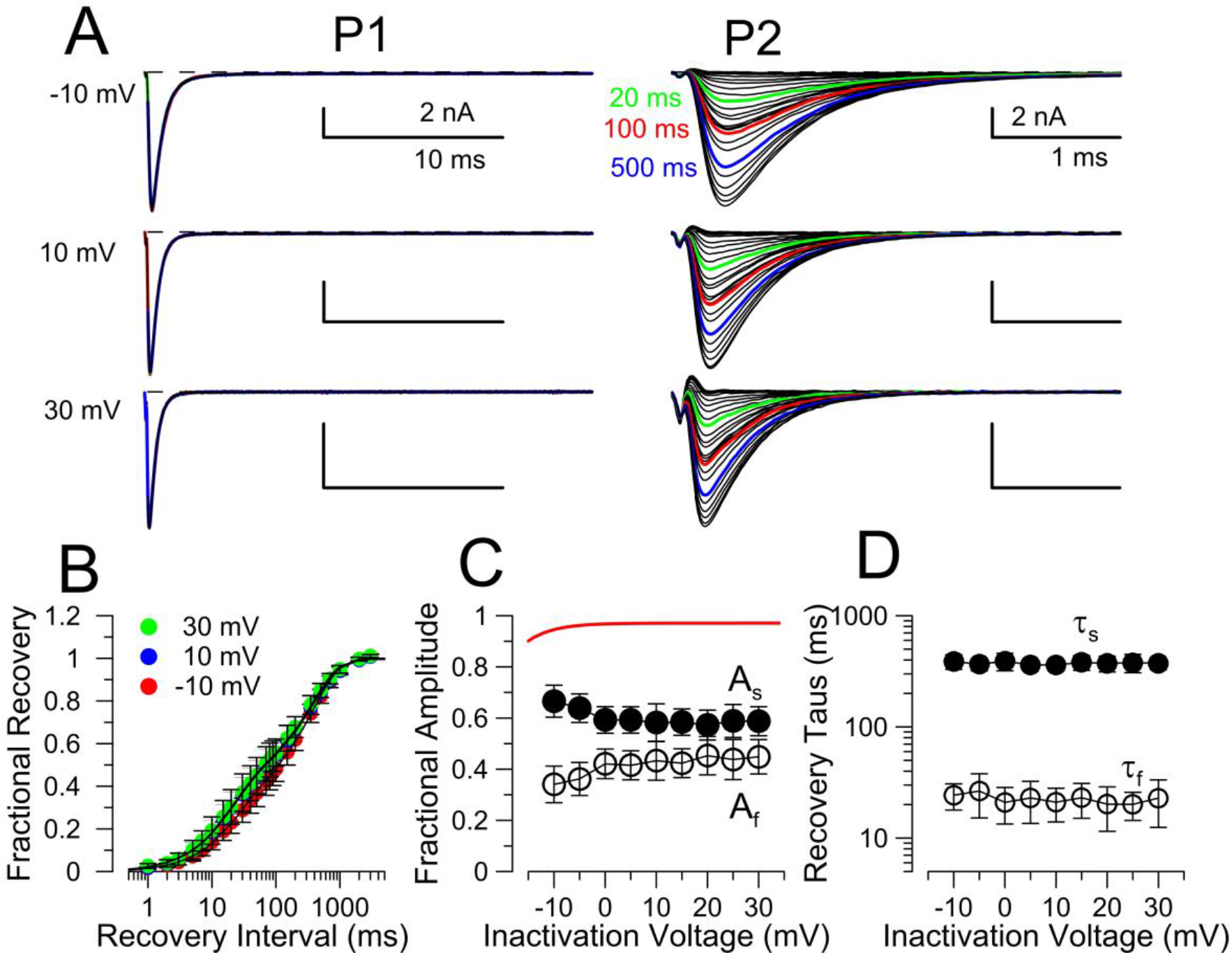
Relative entry into slow and fast recovery pathways is voltage-independent above −10 mV. (A) The standard PP protocol was employed with a 25 ms inactivation depolarization (P1) to voltages from −10 mV to +30 mV, with recovery intervals (at −80 mV) from 1 ms to 3 s, followed by a final 5 msec test step (P2) to the initial inactivation voltage. Traces show currents evoked during P1 on the left for a given voltage, and then during P2 on the right. From top to bottom, traces reflect inactivation voltages of −10, 10, and +30 mV. Green, red, and blue traces show currents following recovery intervals of 20, 100, and 500 ms, respectively. Over this range of voltage, activation of Nav conductance is near maximum (Fig. 1C). (B) Averaged recoveries following inactivation at the indicated voltages (−10 mV, n=12 cells; 10 mV, n=13 cells; +30 mV, n=11 cells; mean ± STD with lines reflecting best fits of a double exponential with the following parameters. Following inactivation at −10 mV, A_f_=0.33 ± 0.01, *τ_f_*=23.1±1.3 ms, A_s_=0.67 ± 0.01, τ_s_ = 372.4 ± 14.1 ms; at +10 mV, A_f_=0.42 ± 0.02, *τ_f_*=20.0±1.1 ms, A_s_=0.58 ± 0.01, τ_s_ = 376.1 ± 20.2 ms and, at +30 mV, A_f_=0.42 ± 0.01, *τ_f_*=19.4±1.0 ms, A_s_=0.58 ± 0.01, τ_s_ = 361.4 ± 19.2 ms. Note that the points for recovery at +10 mV are largely obscured by those for recovery at +30 mV. (C) Mean values (±STD) for the amplitudes of the fast (open black circles, Af) and slow (filled black circles, As) recovery components determined from fits to recovery from at least 11 cells at each voltage are plotted as a function of different inactivation voltages. The red line corresponds to the Nav GV curve from Fig. 1C. (D) Mean values (±STD) for fast and slow recovery time constants (*τ_f_* and τ_s_) for the same set of cells are plotted as a function of the inactivation voltage.

These results indicate that there is little intrinsic differential voltage-dependence of entry into inactivated states, i.e, relative amplitude of fast and slow recovery remain constant, and, furthermore, rates of recovery are not affected by the voltage at which inactivation occurred. Although this protocol *per se* does not fully distinguish between a Scheme 1 or Scheme 2 type of inactivation, the results suggest that changes in voltage do not alter the distribution among states leading to fast and slow recovery.

To further test whether any manipulations might alter the distribution of channels among different recovery pathways, we utilized a protocol in which recovery following inactivation produced by a 5 msec step to 0 mV was examined either with or without a subsequent 50 msec depolarization to +70 mV just preceding the recovery period (Fig. 6A,B). Similar to the impact of increasing duration of depolarizations (Fig. 4), once inactivation has occurred, an additional 50 ms depolarization to +70 mV produces only a modest reduction in the fraction of channels that recover through the fast pathway (Fig. 6C-E). Any additional reduction in the fraction of channels that recover through the fast pathway is generally consistent with the reduction that is observed for increases in inactivation step duration from 5 ms to 55 ms at 0 mV (Fig. 4G). Thus, once inactivation has occurred, despite an additional 50 ms at a more strongly depolarized potential, there is little change in the relative ratio of channels in states leading to slow recovery pathways relative to channels in states leading to fast recovery.

**Figure 6.**
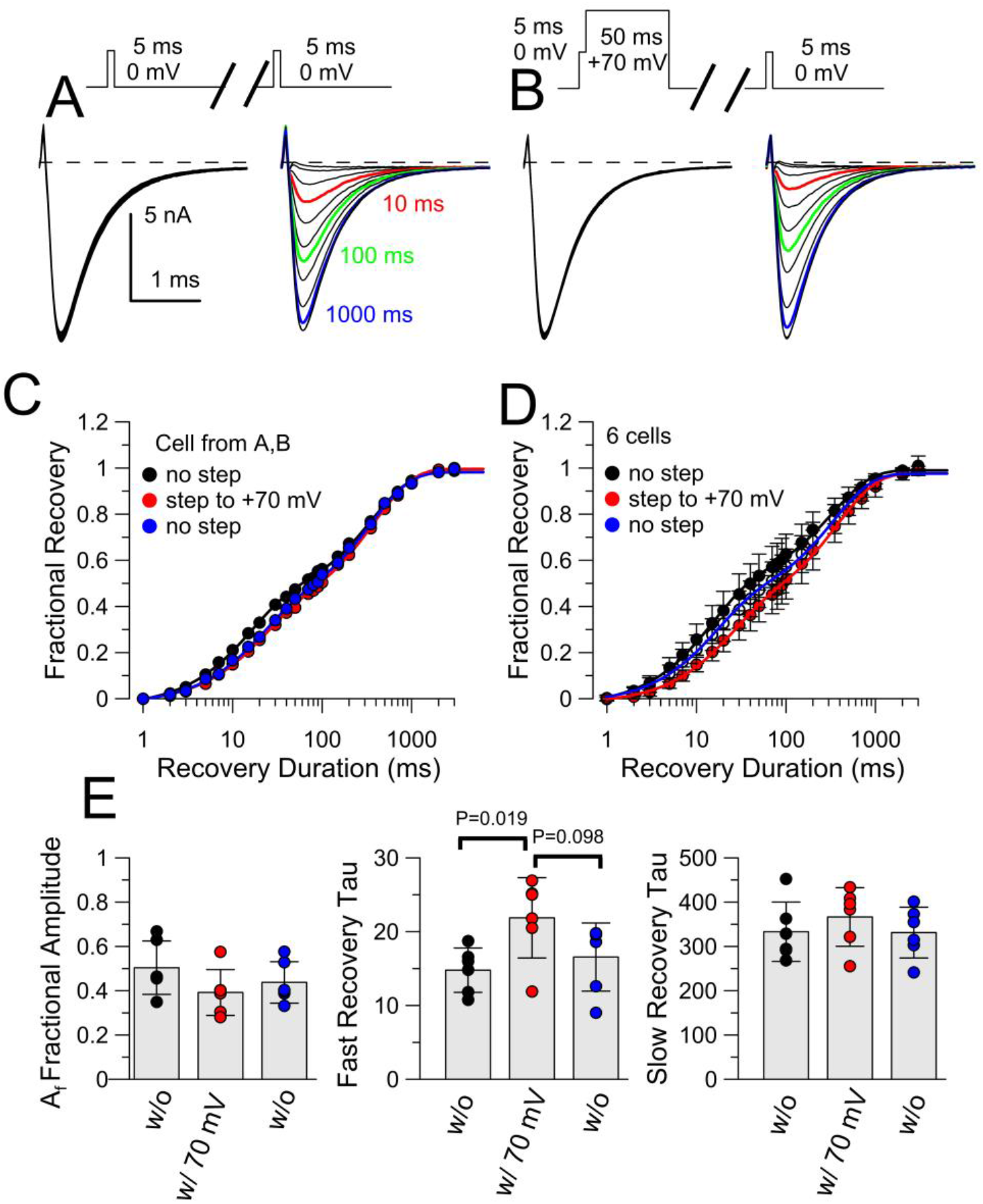
Once inactivation has occurred, additional prolonged depolarization produces only minor changes in distribution between fast and slow recovery. (A) A standard paired pulse protocol (5 ms to ±0 mV) was used to examine recovery from inactivation at −80 mV with recovery in response to the second pulse shown on the right. (B) Following inactivation at ±0 mV, a 50 ms step to ±70 mV preceded the recovery step to −80 mV, with individual recovery tests on the right. (C) Fractional recoveries for the cell shown in A and B are plotted, along with an additional control protocol without the ±70 mV step. Lines are fits of a double exponential function to the recovery time courses. With no step to ±70 mV, Af =0.46±0.01 (mean ± c.l.), *τ_f_*-=14.8 ± 0.7 ms, A_s_=0.56±0.01, and τ_s_=363.0 ± 14.6 ms; with a step to ±70 mV, A_f_=0.39±0.01, τ_f_=20.5±1.2 ms, A_s_=0.63±0.01, and τ_s_ = 388.7 ± 16.6 ms; after return to a protocol with no step to ±70 mV, A_f_=0.40±0.01, *τ_f_*=19.6±1.4 ms, A_s_=0.6-±0.01, and τ_s_ = 354.9 ± 19.2 ms. (D) Average recovery time courses are shown for 6 cells for the indicated conditions. Initial control recovery after a 5 ms step to 0 mV: A_f_=0.51±0.02, *τ_f_*=14.1±1.0 ms, A_s_=0.51±0.01, and τ_s_ = 328.1 ± 25.5 ms. Recovery following both the 5 ms inactivation step and the 50 ms step to ±70 mV: A_f_=0.40±0.01, *τ_f_*=22.3±2.1 ms, A_s_=0.60±0.01, and τ_s_ = 377.3 ± 25.8 ms. A repeat of recovery following just the 5 ms inactivation step to 0 mV: A_f_=0.44±0.02, *τ_f_*=15.6±1.3 ms, A_s_=0.56±0.01, and τ_s_ = 337.9 ± 24.0 ms. (E) Mean and individuals values for Af, *τ_f_*, and τ_s_ are shown for the set of 6 cells from panel D. There are small differences in Af and tf observed with the ±70 mV step, but similar to that which would expected from a 55 ms inactivation step to 0 mV (see Fig. 4). All error bars in D-E are STD. Except as indicated in the middle panel, all T-TEST comparisons yielded P > 0.1.

Based on the idea that traditional slow inactivation involves slow entry from fast inactivated states (Scheme 1), the results in Figures 4-6 seem more compatible with a behavior involving entry into two separable fast inactivation pathways (Scheme 2). Scheme 1 could only be applicable if rates of conversion between fast and slow recovery states were very rapid at positive voltages. This seems unlikely, since at least over voltages from −120 to −50 mV the results above indicate that exit from the slow recovery pathway ranges from 60-500 ms (Fig. 3C) over voltages from −120 to −50 mV.

### Accumulation of channels in slow recovery paths during repetitive inactivation steps requires two independent inactivation pathways

A prediction that arises from the idea that there are two independent, non-equilibrating recovery pathways for CC Nav current inactivation is that sequential inactivation steps with short recovery times should produce a use-dependent accumulation of channels in the slower recovering inactivated states. Specifically, if brief recovery periods are interposed between inactivating command steps, each recovery period would allow more channels to recover through fast pathways than through the slower pathway. If the two pathways are largely independent, each sequential inactivation step will cause a similar fraction of the available channels to enter the sets of states leading to either slower or fast recovery. Since a short recovery period allows only those channels recovering through fast pathways to become available, a series of inactivating steps will result in the accumulation of channels in the set of states leading to slow recovery, but only if the pathways are separate. In contrast, if at strong depolarizations there is rapid equilibration of channels among states leading to slow and fast recovery pathways, accumulation of channels in slow recovery pathways will not occur. We tested these alternatives in Figure 7. We produced inactivation with a 5 ms to +0 mV which approximates the width of a naturally occurring CC action potential at 0 mV (see Fig. 10). We compared recovery from inactivation following a single inactivating step (Fig. 7A,D) to 0 mV to that after trains of five (Fig. 7B), ten, or twenty (Fig. 7C) inactivation steps, with 15 ms between the start of each step (effective frequency of 66.6 Hz). After a train of five inactivating steps (Fig. 7B), there is marked diminution in amplitude of the P2 response during recovery durations of 100 ms and less, with full recovery occurring by 3 sec. With a train of twenty inactivating steps (Fig. 7C), there is only a small additional decrement in P2 currents at short recovery durations over that seen with a train of five steps. Recoveries from the cell shown in 7A,D) are plotted in Figure 7E with compiled data (N=19 cells) in Figure 7F. Overall, the fraction of recovery through fast pathways decreases from 0.5 ± 0.06 (+STD; N=19) following a single inactivation step to 0.27 + 0.06 (same 19 cells) with a 20 pulse train (Fig. 7G). In addition to the changes in amplitude ratios, the fast time constants exhibited some slowing as a function of number of pulses in a train (Fig. 7H), while the slow time constants exhibited no change except for a small difference observed with the 20 pulse train (Fig.7H). Overall, these results demonstrate that rat Nav currents exhibit a pronounced use-dependence in availability that arises from the differential recovery between the two separate inactivation pathways. It is important to realize that the five pulse train of 5 ms steps (total of 25 ms depolarization) produces accumulation in slow recovering states, while increasing a single 5 ms step to 25 ms has very little effect on the distribution between slow and fast recovering states (as in Fig. 4). Thus, during repetitive depolarizations, differential recovery between fast and slow pathways allows use-dependent accumulation of channels in slowly recovering states.

**Figure 7.**
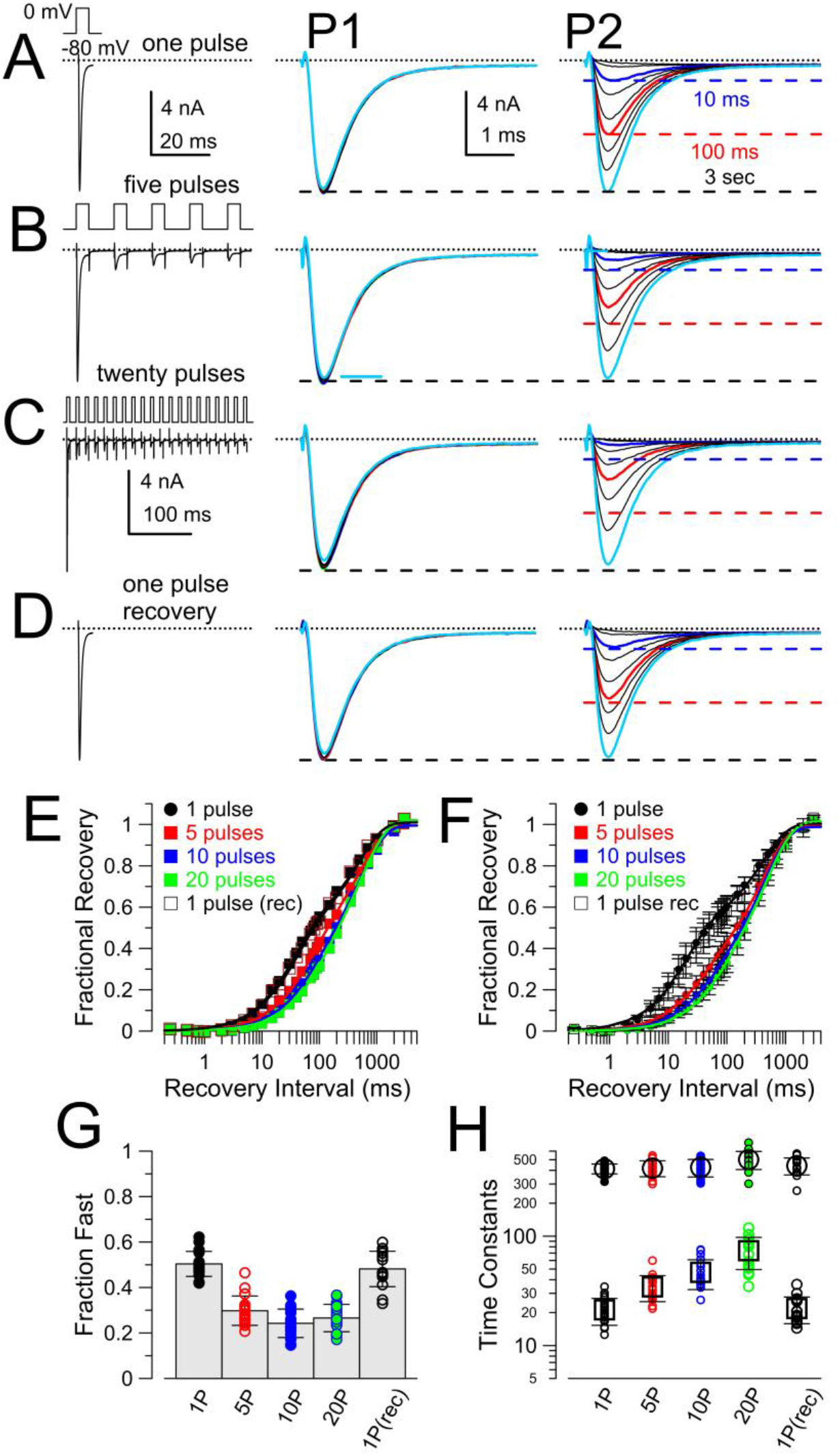
Repetitive stimuli result in use-dependent accumulation of Nav channels in slow recovering states. (A) The standard PP protocol was used to elicit Nav current with a 5 ms step to 0 mV (P1), with recovery intervals at −80 mV from 0.3 ms to 3 sec preceding another 5 ms step (P2) to 0 mV. P1 currents are on the left, while P2 currents on the right followed recovery durations of 1, 3, 10, 20, 50, 100, 200, 500, and 3000 ms. Horizontal colored lines and colored traces highlight traces following 10, 100, and 3000 ms. (B) A train of five 5 ms steps to 0 mV was used as the P1 stimulus, with 10 ms intervals between steps to 0. Currents on the right are as in A. Traces at 10 (blue) and 100 (red) ms exhibit marked diminution compared to that in panel A. Dotted lines reflect the amplitudes of the recoveries for 10, 100, and 3000 ms from A. (C) Traces are as in panel B, but for a train of twenty 5 ms steps to 0 mV, with 10 ms between each step in the train. Dotted lines on the right again correspond to fractional recoveries observed in panel A for 10, 100, and 3000 ms. (D) Following stimulation with the trains, the standard single inactivation pulse was again applied, as in A, showing that, following a single inactivation step, fast recovery returns to its initial amplitude. (E) Fractional recovery from inactivation is plotted for the cell in A-D, with each curve fit with a two exponential function with the following values: for initial control recovery, A_f_=0.49±0.03,*τ_f_*=34.5±3.0 ms, A_s_=0.52±0.02, and τ_s_=481.3±52.5; following the five-pulse train, A_f_=0.40±0.03, *τ_f_*=59.7±6.0 ms, A_s_=0.60±0.03, τ_s_=515.6±46.1 ms; following a ten-pulse train, A_f_=0.31±0.05, *τ_f_*=68.4±12.4 ms, A_s_=0.68±0.05, τ_s_=534.8±58.3 ms; following a twenty-pulse train, A_f_=0.29±0.04, *τ_f_*=84.0±12.5 ms, A_s_=0.73±0.04, τ_s_=562.8±44.0 ms; after recovery, A_f_=0.47±0.02, *τ_f_*=36.2±3.2 ms, A_s_=0.54±0.23, τ_s_=475.3±48.5 ms. (F) Averaged values (±STD) for fractional recovery are plotted for a set of 19 cells (recovery runs were only obtained for 16 of the cells). For the initial control protocol, recovery parameters (mean±90% c.l.) are A_f_=0.50±0.02, *τ_f_*=19.6±1.4 ms, A_s_=0.51±0.02, τ_s_=404.3±39.4 ms; for the five-pulse P1 stimulus, A_f_=0.30±0.02, *τ_f_*=33.6±3.3 ms, A_s_=0.70±0.02, τ_s_=413.6±22.9 ms; for the ten-pulse P1 stimulus, A_f_=0.24±0.03, *τ_f_*=45.2±6.4 ms, A_s_=0.75±0.02, τ_s_=418.6±24.3 ms; for the twenty-pulse P1 stimulus, A_f_=0.26±0.04, *τ_f_*=67.6±12.0 ms, A_s_=0.76±0.04, τ_s_=493.3±38.0 ms; following return to the single pulse protocol, A_f_=0.48±0.02, τ_f_=20.5±1.7 ms, A_s_=0.53±0.02, τ_s_=436.3±44.4 ms. (G) Mean values (±STD) of the fast recovery amplitude for each condition are plotted along with the best fit values from each individual cell in the set of 19 cells. Control (1P) and recovery (1P(rec)) fast amplitude differed from all other protocols (5P, 10P, 20P) at P=0.000 (KS Test). 1P v. 1P(rec): 0.085; 5P v 10P: 0.049; 5P v. 20P: 0.462; 10P v. 20P: 0.742. (H) Fast and slow time constants (±STD) are plotted for each of the recovery protocols, along with the individual determinations for each cell (small symbols). K-S P values for comparisons of fast time constant values were: for 1P v. 5P, 0.000; 1P v. 10P, 0.000; 1P v. 20P, 0.000; 1P vs 1P(rec), 0.957; 5P v. 10P, 0.018, 5P v. 20P, 0.000, 10P v. 20P, 0.002; 20P v. 1P(rec), 0.000. For comparisons of slow time constants: 1P differed from 5P, 10P, and 20P at 0.000; 1P v. 1P(rec), 0.354; 5P v. 10P, 0.116; 10P v. 20P, 0.956; 20P v. 1P(rec), 0.000.

The protocol in Fig. 7 utilized a train frequency much higher than is ever observed in CCs. We therefore compared trains of ten 5 ms steps to 0 mV applied at frequencies of 1, 4, and 10 Hz. 1 Hz is the frequency at which some CCs spontaneously fire (Martinez-Espinosa et al., 2014; Vandael et al., 2015), whereas small depolarizations can evoke firing with instantaneous frequencies up to 10-20 Hz (Solaro et al., 1995). Following inactivation produced either by a single depolarization to 0 mV (Fig. 8A), a 4 Hz train of 10 pulses (Fig. 8B), or a 10 Hz train of 10 depolarizations (Fig. 8C), variable recovery durations were applied prior to define the recovery time course (as shown by P2, the step to 0 mV applied after the train is completed). Qualitatively, increases in train frequency result in reductions in amplitude of currents evoked during P2, but these reductions are most apparent for recovery durations up through 100 ms, whereas by 3 sec virtually all channels have recovered from inactivation. At all tested frequencies, peak Nav current evoked by sequential steps to 0 mV during the train exhibits gradual reduction, with residual peak Nav current during a 66.7 Hz train being less than 0.1 of the initial peak current (Fig. 8D) consistent with persistence of channels in inactivated states even at the lowest train frequencies. The time courses of recovery from inactivation show that fast recovery is reduced as train frequency increases (Fig. 8E), with little difference between recovery from a single pulse or a 1 Hz train of 10 pulses (Fig. 8F). The fraction of channels in fast recovery states appears to diminish as frequency is increased (Fig. 8F). Increasing train frequency also appear to slow the fast component of recovery from inactivation, with little effect on the slow component of recovery (Fig. 8G).

**Fig. 8.**
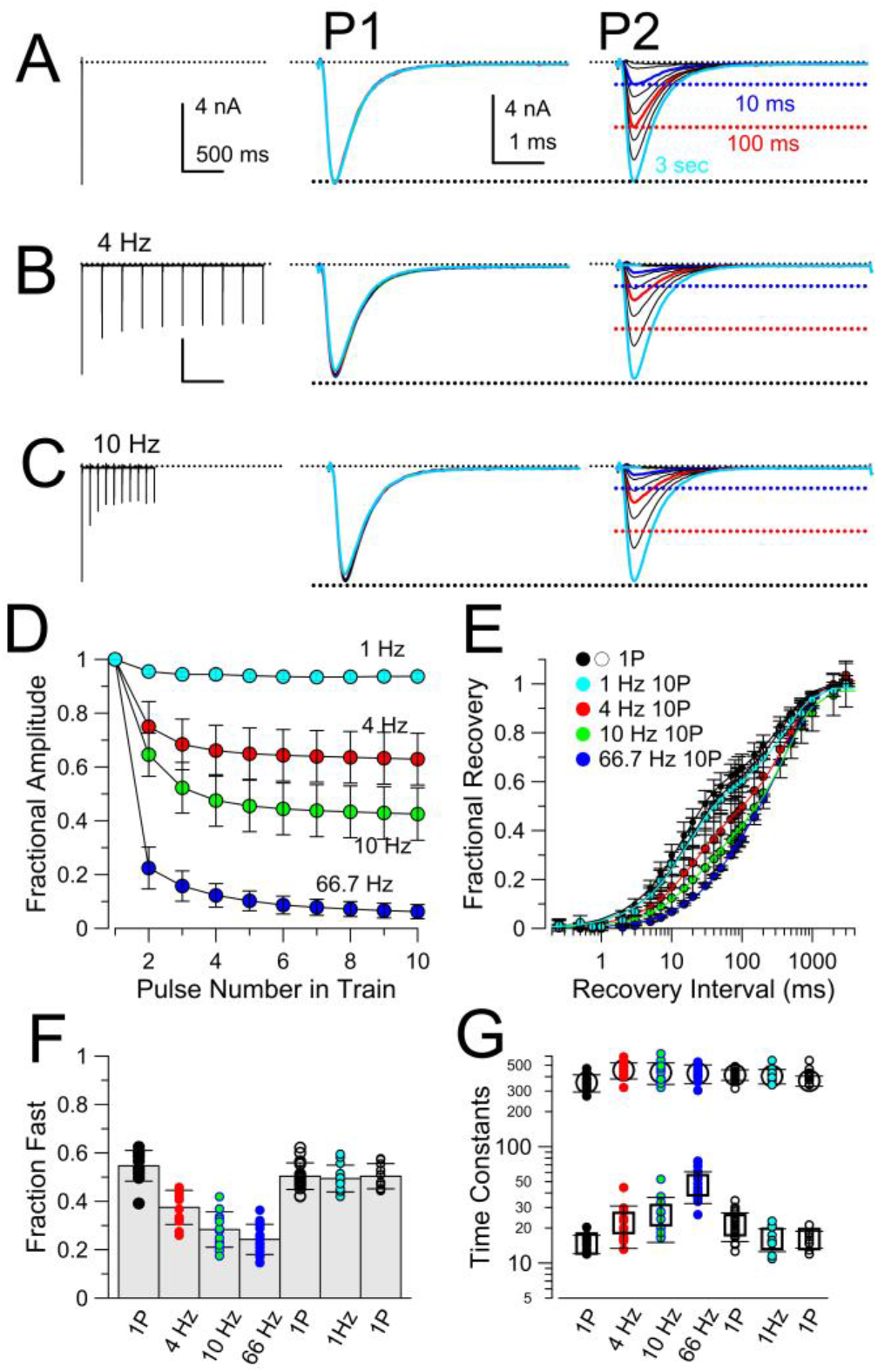
Decrease in peak Nav amplitude during ten pulse trains applied at different frequencies is associated with accumulation of channels in slow recovery pathways. (A) A standard single 5 ms step to 0 mV was used to produce inactivation (P1), with recovery at −80 mV preceding a test step (P2) to 0 mV. Dotted lines highlight amplitude following 10, 100 and 3000 ms of recovery. (B) From the same cell as in panel A, a 4 Hz, ten pulse train of 5 ms steps to 0 mV was applied prior to the standard recovery step to −80 mV with dotted lines reflecting recovery amplitudes in A. (C) From the same cell as in A-B, a 10 Hz train preceded the recovery steps. (D) The decrement in peak Nav current amplitude is plotted for 1, 4, 10, and 66.7 Hz trains of 10 pulses applied to 0 mV. Means±STD.(E) Time course of recovery from inactivation following different stimulus trains. 12 cells were used for 4 and 10 Hz trains, which also included a control set of recovery determinations with the standard single pulse (1P) protocol. 13 different cells were used for the 1 Hz protocol, which were also tested with the 1P protocol. 19 cells (same as in Fig. 7) were used with the 66.7 Hz 10 pulse protocol, which also included the 1P protocol as a control. Fit values were for 1P, 4Hz, and 10 Hz cells were as follows. For 1P: A_f_=0.55±0.02, *τ_f_*=14.7±0.8 ms, A_s_=0.44±0.02, τ_s_=354.3±17.2 ms; for 4 Hz, A_f_=0.37±0.02, τ_f_=22.1±2.5 ms, A_s_=0.64±0.02, τ_s_=453.2±20.7 ms; for 10 Hz, A_f_=0.28±0.02, *τ_f_*=25.8±3.1 ms, A-_s_=0.69±0.04, τ_s_=433.7±26.3 ms. For 1P and 1 Hz cells, 1P: A_f_=0.50±0.02, *τ_f_*=16.1±1.0 ms, A-_s_=0.50±0.02, τ_s_=367.3±13.0 ms; for 1 Hz, A_f_=0.49±0.02, *τ_f_*=16.1±1.3 ms, A_s_=0.49±0.02, τ_s_=403.2±20.3 ms. For 1P and 66.7 Hz cells, 1P: A_f_=0.50±0.02, *τ_f_*=19.6±1.9 ms, A_s_=0.51±0.02, τ_s_=404.3±51.6 ms; for 66.7 Hz, A_f_=0.24±0.03, *τ_f_*=45.2±8.6 ms, A_s_=0.75±0.03, τ_s_=493.3±49.9 ms. (F) Means, standard errors, and individual values for the fitted fast component amplitude for all individual cells for the indicated stimulus protocols. KS-test P values for 12 cells compared wth 1P, 4 Hz, and 10 Hz protocols: 1P vs. 4 Hz, 0.000; 1P vs. 10 Hz, 0.000; 4 Hz vs. 10 Hz, 0.066. For 13 cells compared with 1P and 1 Hz protocols: 1P vs. 1 Hz, 0.881. For 19 cells, compared with 1P and 66.7 Hz protocols: 1P vs. 66.7 Hz, 0.000. (G) Means (±STD) and individual determinations of slow and fast time constants are displayed for the indicated conditions. For cells tested with 1P, 4 Hz, and 10 Hz protocols, KS P values, for the fast recovery time constant, were 0.019 (1P v. 4 Hz), 0.001 (1P v. 10 Hz), and 0.433 (4 Hz v. 10 Hz). For the 1P vs. 1 Hz train comparison, P = 0.638. For the 1P vs. 66.7 Hz train comparison, P=0.000. For slow time constants, for 1P v. 4 Hz, P=0.001; for 1P v. 10 Hz, P=0.186; for 4 Hz v. 10 Hz, P=0.186. For the 1P vs. 1 Hz train comparison, P=0.341. for the 1P vs. 66.7 Hz comparison, P=0.462.

The above results generally support the idea that there are two separable fast inactivation pathways with differential recovery properties. However, two aspects of the results seem inconsistent with the simplest version of a model with two independent, separable pathways. First, the slowing of the fast recovery time constant is not consistent with the most simple model of two independent and competing inactivation and recovery mechanisms. Second, during the repetitive pulse train, what limits the fractional accumulation in the slow recovery pathways to about 0.75? Might the inability to drive channels fully into slow recovery pathways reflect the possibility that some channels may not contain the fast inactivation/slow recovery machinery. Although we have no explanation for the apparent slowing of the fast component of inactivation as train frequency (Fig 8G) or train duration (Fig. 7H) is increased, we sought to address the second question by asking whether a simple approximation of the simple dual-pathway inactivation model could be consistent with the observations. To accomplish this, we used initial fractional availability before a train based on Fig. 1E, and time constants of recovery from inactivation from Fig. 3C. Furthermore, we assume that a 5 ms depolarization to 0 mV fully inactivates all available channels, and half go into slow recovering pathways and half into fast. Based on this, we calculated the fractional occupancy of channels in closed states, slow recovery inactivated states, and fast recovery inactivated states for times immediately preceding each depolarization in the train, and then immediately after each depolarization in the train (Fig. S1A). From the fraction of channels in closed states prior to each depolarization, the predicted run-down in peak Nav current was determined (Fig. S1B-C) and matched very well with the measured decreases in Nav current with different train frequencies (Fig. 8D). Plots of the fractional occupancy of channels in slow and fast recovery pathways show that at 1, 4, and 10 Hz, essentially all channels in fast recovery pathways recover during the 995, 245, and 95 ms recovery intervals (Fig. S1D), respectively. However, at 66.7 Hz, the 10 ms recovery interval is insufficient to allow full recovery from fast inactivation. The slow decrease in fraction of channels in fast recovery pathways reflects the slow increase in channels in slow recovery pathways. The state occupancies immediately after termination of the 5 ms depolarization better reflect what would be expected for the fraction of fast and slow recovery components during recovery from inactivation (Fig. S1E-F). For the simplest case, we assumed that, irrespective of train frequency, recovery time constants were identical (Fig S1E). In this case, as train frequency increases, the fraction of channels in fast recovery pathways monotonically decreases (Fig. S1G), with a value of about 0.1 after 10 pulses. This would seem to deviate from the experimentally observed tendency of the fast component to reach a limit of about 0.25 fractional availability. However, the results also indicate that the apparent fast time constant after a 10 pulse train is slowed at higher train frequencies (Fig. 8F). We therefore modelled expectations for fast and slow components with the assumption that the fast time constant slows with train frequency (Fig. S1F). Although slowing the fast time constant of recovery has minimal impact on the fractions of slow and fast components at 1, 4, and 10 Hz, the value of the fast time constant can impact substantially on the overall rate of accumulation in slow recovery pathways and also impact on the fractional occupancies in slow and fast pathways after long trains (Fig. S1F-G). Thus, based on experimentally measured values, this simple approach can generally recapitulate the frequency-dependence reductions in Nav current amplitude and also the general properties of the frequency-dependence of changes in the fraction of fast recovery channels (Fig. S1G). Based on these considerations, we think that various protocols never drive more than about a fraction of 0.75 into slow recovery pathways simply reflects the relative time constants of slow and fast recovery at higher stimulus frequencies, and not a population of channels lacking the slow recovery mechanism.

The above protocols have focused on protocols that define properties of the dual-pathway fast inactivation following depolarizations that produce full activation and during conditions where recovery to resting states predominates. However, the normal resting potential of CCs, typically considered between −45 and −55 mV (Martinez-Espinosa et al., 2014; Vandael et al., 2015), spans a range where there is likely to be considerable equilibration of channels in and out of closed, inactivated, and even, to some extent, open states. Therefore, in the sections below we examine protocols that evaluate recovery from inactivation following inactivation over voltages in the range of −40 to −70 mV and similarly test onset of inactivation over this same voltage range. These tests only qualitatively describe the extent to which channels may occupy slow and faster recovery pathways over membrane potentials at which CCs spend most of their time between APs.

### Nav current availability is markedly reduced by repetitive AP clamp waveforms

Results above indicate that occupancy of channels in slow and fast recovery pathways near normal resting potentials will affect Nav availability. Here, to assess the impact of holding potential and firing frequency on Nav availability under conditions more reflective of physiological circumstances, we have used AP-like waveforms to examine changes in Nav availability during repetitive activity. To accomplish this, a spontaneously occurring AP was recorded from a rat CC and then used as a voltage-clamp command. The AP waveform included 5 ms at −52 mV before the upswing of the AP and then a period of about 35 ms corresponding to an afterhyperpolarization (AHP). Experiments were done on dissociated CCs maintained in short-term culture which allowed the use of perforated-patch recording methods, with 20 mM cytosolic Cs^+^ to inhibit outward current. AP clamp waveforms were applied at frequencies of either 4, 10, or 20 Hz at holding potentials between the AP waveform. With 120 mM extracellular Na^+^ and 2 mM extracellular Ca^2+^, the currents evoked by the AP clamp waveform exhibit a biphasic inward current (Fig. 9A), with strong inward current activated during the rising phase of the AP command waveform, essentially 0 net inward current at the AP peak, and then a secondary reduced peak of inward current during AP repolarization. The amplitude of the initial inward current during the AP rising phase exhibits strong suppression at a 10 Hz AP frequency, while the amplitude of the secondary inward current is unaltered at 10 Hz. This indicates that essentially all Nav current is fully inactivated at the time of the secondary phase of inward current during the first AP in a train and that the secondary inward current likely arises exclusively from voltage-dependent Ca^2+^ (Cav) current. Using this approach, the impact of different holding potentials (−50, −60, −70, and −80 mV) both before and between the AP waveforms was examined (Fig. 9B). As expected from the steady-state inactivation behavior of the Nav current, the maximum peak inward current activated by the AP clamp waveform varied substantially over holding potentials from −50 mV through −80 mV. Furthermore, as shown for AP waveforms applied at 10 Hz, at all recovery potentials there was marked diminution of the AP waveform evoked Nav current (Fig. 9B).

**Figure 9.**
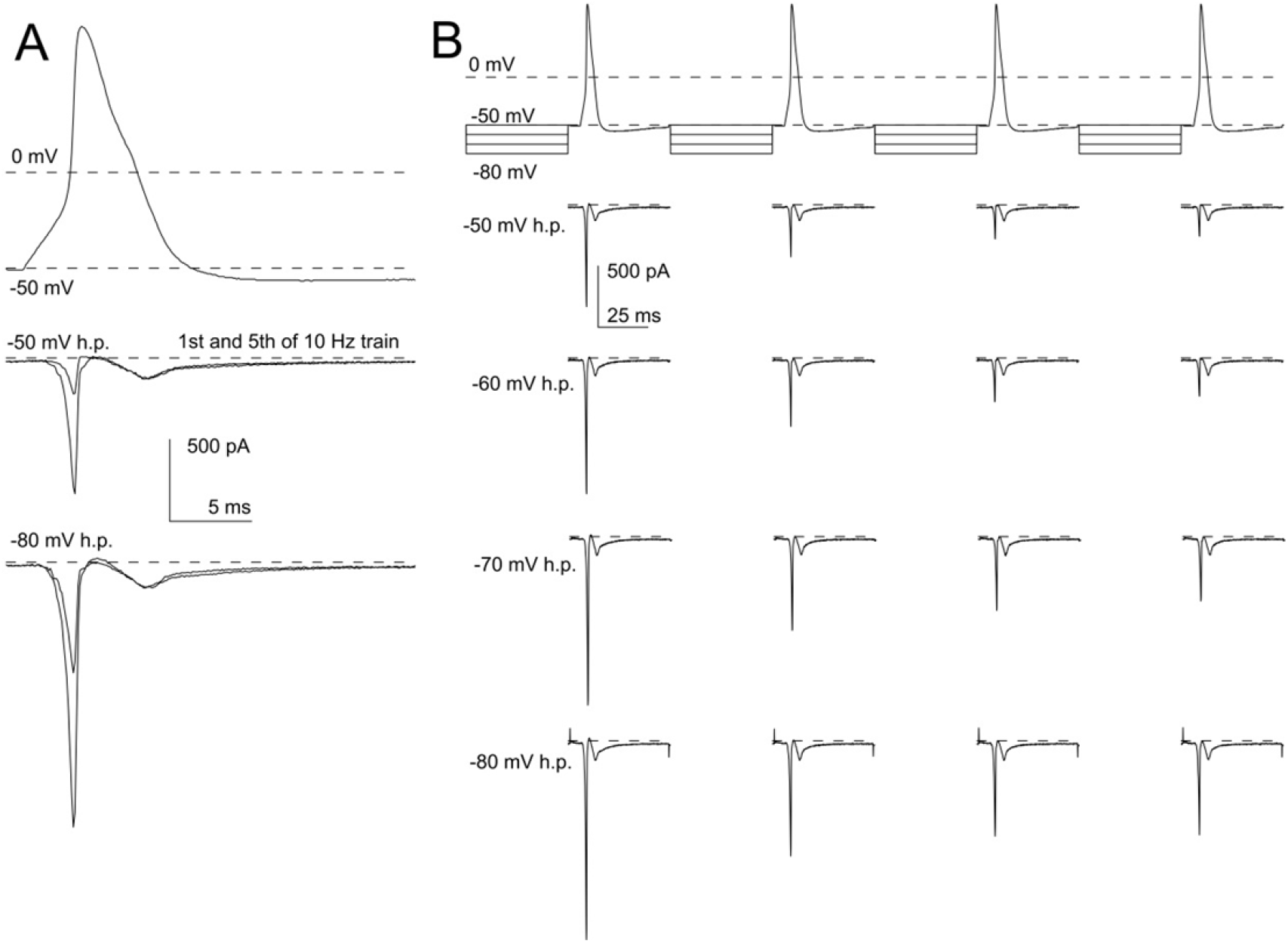
Inactivation of Nav current by AP clamp waveforms. (A) Top trace shows an AP recorded from a rat CC with the perforated patch-method with normal Na^±^ and K^±^ gradients. The AP waveform was then used as a voltage-clamp waveform for all traces in A and B. For a given train of AP clamp waveforms, a cell was held at different holding potentials, −50 through −80 mV, before and between each AP clamp waveform. For the example in A, 100 ms separated the initiation of each individual AP clamp waveform. The traces below the AP waveform show currents activated from a holding potential of −50 mV or −80 mV (bottom), for the 1^st^ and 5^th^ evoked current in a 10 Hz train. Note that the inward current during the falling phase of the AP command waveform does not change in amplitude between the 1^st^ and 5^th^ waveform, reflecting Cav current, while the early inward current shows marked diminution, indicative of Nav current inativation. (B) The full command waveforms for the first four APs in a 10 pulse train are shown along with the currents evoked by the AP command waveforms for holding potentials from −50 mV through −80 mV.

When the AP-evoked peak inward current amplitudes are plotted as a function of the overall elapsed time of the train of AP commands (Fig. 10A-C), the initial peak inward current is reduced as holding potential is made more positive. Irrespective of train frequency, a gradual reduction in Nav current amplitude occurs over the first 5-6 APs, largely reaching a plateau after that (Fig. 10A-C). At −50 mV holding potential, the peak inward current can be reduced at a 10 Hz AP frequency to less than 20% of that available from a −80 mV holding potential (Fig. 10B). Even at a 4 Hz frequency, the peak Nav current amplitude is reduced to around 20% of the peak available from a −80 mV holding potential and less than 50% of that available at a −50 mV holding potential. Overall, this experiment indicates that, during normal AP frequencies and at reasonable membrane potentials, Na^+^ channel availability is quickly reduced to about 10-25% of the full availability, with essentially all Nav channels being inactivated during a single AP waveform.

**Figure 10.**
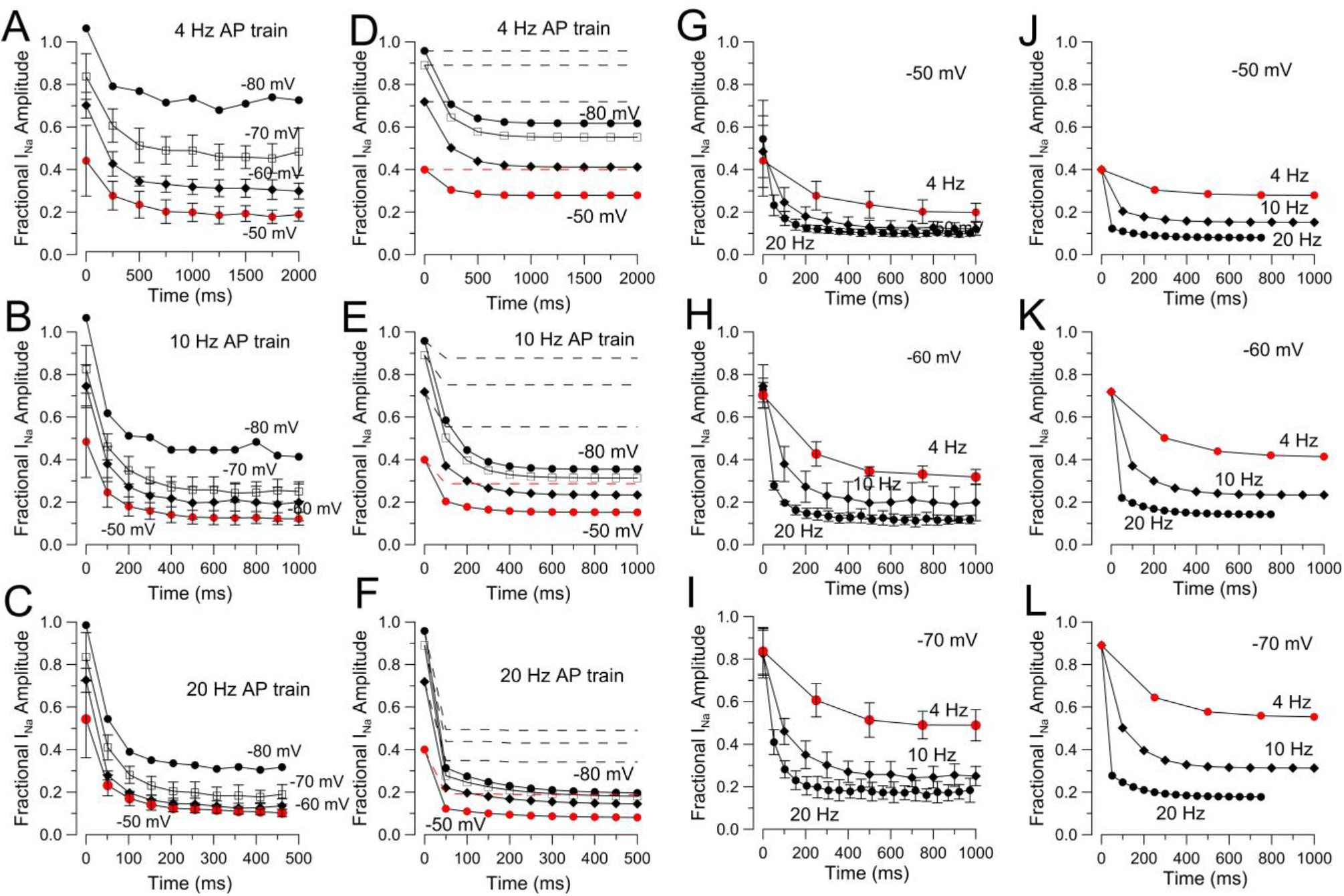
Diminution of AP-evoked peak inward current with different holding potentials and AP frequencies. (A) Peak inward current evoked by each AP clamp waveform applied at 4 Hz was normalized to the maximal available current defined in a given cell using a standard Nav activation protocol (Fig. 1A) from a holding potential of −80 mV. At −50 mV the initial AP is reduced in amplitude by about 50% compared to that from −80 mV and diminishes to less than 20% by the 5^th^ AP command. The time base corresponds to the total elapsed time of the protocol as shown in Figure 11B. Number of cells: −50 mV. N=4; −60 mV, N=3; −70 mV, N=2; −80 mV, N=1. Error bars here and in other panels are STD. (B) Normalized inward current amplitude at different resting/holding potentials is shown for a 10 Hz AP train. (C) Normalized inward current amplitude is shown for a 20 Hz AP train. (D) Predicted decrements in peak Nav current were calculated as following. Availability at a given holding potential was determined from the steady-state inactivation curve (Fig. 1E) with a 250 ms prepulse. Measured fast and slow recovery time constants (Fig. 3C) were used to calculate fractional recovery during given sojourns at particular voltages. The predicted decrement based on the protocol of Fig. 10B was calculated for the 4 Hz AP train assuming that all Nav current inactivates during each AP, with the first AP driving half the channels into a slow recovery pathway and half into a fast recovery pathway. Recovery than occurs following the AP for a period of 36 msec corresponding to an AHP ranging from −56 to −50 mV; for simplicity, we assumed −50 mV for this interval. Then, the additional recovery that occurs during the interval between sweeps at the specified recovery voltages of −50, 60, −70, or −80 mV was calculated. For comparison, the calculated AP decrement when all inactivation is exclusively via a fast recovery pathway is also shown (dotted lines, red dotted line indicates −50 mV). (E) The same calculations as in D were done for a 10 Hz train showing the deeper decrement in AP amplitudes, because of slower recovery intervals. (F) Fractional decrement in AP amplitude calculated for the 20 Hz train is shown. (G) The data from panel A-C are recast to directly compare 4, 10, and 20 Hz trains all for a given holding potential (−50 mV). (H) As in G, but for −60 mV. (I) As in G, but for −70 mV. (J-L) The calculated decrements for the conditions plotted in G-I are shown.

To evaluate whether these decrements in AP evoked currents might be consistent with the measured properties of dual-pathway inactivation, we followed the general procedures outlines for Fig. S1 in which we determined the predicted decrement in Nav current amplitude during application of trains of 5 ms depolarization, taking into account the specific details of the AP train waveforms (Fig. 9B). Using measured time constants of recovery from inactivation at different voltages (Fig. 3C), the calculated decrements in evoked current amplitude during a train of ten AP clamp waveforms (Fig. 10D-F) closely follow the measured decreases (Fig. 10A-C).

Furthermore, we also tested removal of slow recovery from inactivation, with the assumption that all inactivation is by a single fast inactivation pathway (dotted lines in Fig. 10D-F). With 10 Hz and 20 Hz trains, an initial decrement in peak Nav current amplitude is predicted, but this is complete within 2-3 APs, consistent with the absence of any slow accumulation of channels in slow recovering inactivated states. We also recast the results and analysis in panels A-F to better focus on comparisons among different train frequencies at a single holding potential. The correspondence of the measured changes in Nav current amplitude (Fig. 10G-I) with the calculated changes (Fig. 10J-L) based on the two separable inactivation pathways idea provides further support for this idea and illustrates the importance of the slow recovery pathways in defining changes in Nav availability during AP trains in CCs. When the fractional occupancies of channels in available, slow recovery inactivated, and fast recovery inactivated states are determined (Fig. S2), the analysis shows clearly that occupancy of channels in slow recovery states is the major determinant of the reduction in AP-evoked Nav current amplitude.

## Discussion

The present results establish that rapid inactivation of Nav current in rat CCs involves entry into two distinct pathways of fast inactivation, each associated with differential rates of recovery from inactivation. Our results show that the slow recovery process described here is quite distinct from traditional slow Nav inactivation, as summarized below. Here, we also consider the potential implications of this dual pathway inactivation for AP firing in CCs. We note that, although the variety of protocols used here to probe the contributions of the two pathways, might seem suitable for evaluation of gating models, until the molecular determinants and potential interactions between the pathways are directly tested, we feel any attempt to explicitly model the present observations is premature.

### Slow recovery of Nav channels from inactivation

Beginning with the initial studies of voltage-dependent Nav currents, slow components of recovery from inactivation have been observed in a number of preparations, including squid giant axon (Chandler and Meves, 1970; Rudy, 1978), *Myxicola* axons (Rudy, 1981) and rat sympathetic neurons (Belluzzi and Sacchi, 1986). Typically, such slow recovery from inactivation is a process that also develops slowly, with increases in the duration of the inactivation step leading to increases in the fraction of current that recovers slowly (Silva, 2014). Such slow inactivation behavior, which is thought to involve coupling of slow inactivation to fast inactivation, modeled as in Scheme 1, has been proposed to apply to the cardiac Nav channel, Nav1.5 (Zhang et al., 2013) and other cases (Silva, 2014). Essentially, because slow inactivation is preceded by entry into fast inactivated states, the sojourn of time in fast inactivated states results in accumulation in slow inactivated states.

In contrast to traditional slow inactivation, the results here argue strongly for two functionally distinct competing fast inactivation pathways, with one pathway having slower recovery kinetics. Perhaps the first proposal for two competing inactivation pathways with different recovery kinetics was that for inactivation of Nav current in *Myxicola* axons (Rudy, 1981). However, in that case the slow recovery was much slower than observed here and appears unlikely to play much role under physiological conditions. More recently, as new information regarding the properties of various mammalian Nav channel variants has appeared, additional examples of Nav currents with slow recovery properties likely to be of physiological importance have appeared (Lou et al., 2005; Goldfarb et al., 2007; Shakkottai et al., 2009; Milescu et al., 2010; Venkatesan et al., 2014; Navarro et al., 2020). One of the most intriguing examples, perhaps relevant to our findings with rat CC Nav current, concerns slow recovery from inactivation of Nav1.6 current observed in cerebellar granule cells (Goldfarb et al., 2007; Dover et al., 2010; Goldfarb, 2012). In this case, a slow component of recovery from inactivation has been attributed to members of a family of cytosolic proteins called intracellular fibroblast growth factor homologous factors (iFGFs, but also termed FHFs) (Olsen et al., 2003; Wittmack et al., 2004). The inactivation behavior of Nav1.6 current, either when coexpressed with particular iFGFs or as the native cerebellar granule cell Nav current, is well-described by a dual-pathway, fast inactivation model similar to that given in Scheme 2, as first proposed by Goldfarb et al. (2007). Specifically, inactivation mediated by certain iFGFs is proposed to compete with the intrinsic fast inactivation particle. As a consequence of the slower recovery from inactivation arising from iFGF-mediated inactivation, accumulation of channels in more slowly recovering states can occur. A similar model has also been used to describe Nav current behavior in dorsal raphe neurons (Milescu et al., 2010). The latter case may be more applicable to the present situation in rat CCs, since the dorsal rat raphe neurons typically fire at frequencies only up to 5-20 Hz, while the cerebellar granule cells fire at frequencies in excess of 50 Hz. Two important points regarding this dual-pathway inactivation process are, first, that rates of entry into both the fast and slow recovering populations are comparable and, second, there is no equilibration between the two pathways, i.e., they are strictly competitive inactivation processes (Goldfarb, 2012). In the case of iFGF-mediated inactivation, it has been proposed that the N-termini of specific isoforms of iFGF can move into a position of occlusion in specific Nav channels (Dover et al., 2010; Goldfarb, 2012; Venkatesan et al., 2014) competing with the ability of the IFM triplet of residues in the DIII/DIV α-helical loop of an Nav α subunit to mediate traditional fast inactivation (Patton et al., 1992; West et al., 1992; O’Leary et al., 1995).

Although the present results do not provide any information after the molecular underpinnings of the slower recovery component of fast inactivation in rat CCs, our experiments do place some limits on the relationship between the two inactivation pathways. First, once inactivation has occurred at potentials of 0 mV and more positive, channels which have entered states leading to fast recovery do not seem to interconvert with channels in slow recovery pathways, over times of 4 to 100 ms. Although we observed some diminution in the relative fraction of the fast component with inactivation pulse duration, the time course of this gradual diminution may be better explained by the slow onset of traditional slow inactivation. Second, the relative fraction of channels entering either slower or faster recovery pathways is largely voltage-independent at voltages above 0 mV. Third, once channels have inactivated, stronger additional depolarizations only weakly influence the distribution of channels among states leading to slow and fast recovery paths. These results are all consistent with the idea that two rapidly entered inactivating pathways exhibit essentially no equilibration between them once inactivation has occurred. Overall, these behaviors are consistent with the general inactivation model outlined in Scheme 2, with the uncertainty that our results provide no information about the extent to which entry into I_SR_ states may occur from closed Nav channels. For iFGF-mediated inactivation, Goldfarb has proposed that the iFGF-dependent inactivation can occur from at least up to two or three of the closed states preceding opening (of the four sequential voltage-sensor movements that lead to opening) (Goldfarb, 2012). The I_SR_ component of Nav current in dorsal raphe neurons was modeled with inactivation only occurring from open Nav channels (Milescu et al., 2010). The extent to which closed-state inactivation may occur from traditional fast inactivation versus iFGF-mediated inactivation may impact importantly on use-dependent changes in channel availability and this topic needs to be addressed in future work.

Overall, the slow component of recovery from inactivation in CCs has several key properties that distinguish it from the traditional slow inactivation (Zhang et al., 2013; Silva, 2014). Specifically, a characteristic of classic slow inactivation is that the fraction of current that recovers through slow recovery pathways increases as a function of the duration of the inactivation step (Zhang et al., 2013; Silva, 2014). This requires a mechanism in which entry into slow inactivated states is coupled to occupancy of channels in fast inactivated states. Thus, entry into the slowly recoverying states is itself very slow. In contrast, for rat CC Nav current, even inactivation steps as brief as 5 ms result in entry into inactivated states from which channels exhibit slow recovery and, importantly, as the duration of the inactivation step is increased, there is little change in the fraction that recover through the slow pathway. The protocols used here unambiguously show that the two recovery pathways are independent and exhibit features completely distinct from classic slow inactivation. As such, the inactivation behavior of Nav current in rat CCs represents a dual-pathway, fast inactivation process, and it is the differential rates of recovery from inactivation of the two pathways that confers physiological importance on the mechanism.

### Potential physiological consequences of dual-pathway, fast inactivation in rat CCs

In rat CCs, a single AP waveform produces almost complete inactivation of Na^+^ channels (Fig. 11A). Following inactivation produced by a single AP, about half the channels will recover through a traditional fast recovery pathway and half through the slower recovery pathway. Once channels have inactivated at very positive voltages, there is no conversion between slow and fast recovery pathways. Thus, between APs occurring at about 5 Hz, most channels that recover from inactivation will be those from the traditional fast inactivation/fast recovery states. With bursts of APs, each subsequent AP will therefore produce accumulation of channels in slow recovery pathways. Even at stimulation frequencies of less than 20 Hz (Fig. 10), about 80% of Na^+^ channels become unavailable for activation at resting potentials of −50 to −70 mV for periods of 100s of ms. The properties of Nav inactivation in rat CCs therefore appear to be well-tuned so that Nav availability can be markedly altered during different patterns of electrical activity. This property may allow Nav channels to participate critically in the determination of the ability of a cell to initiate an AP during mild depolarizations, thereby sculpting cell firing rates.

Based on these considerations, removal of the slow recovery process might help sustain higher frequency firing in CCs. Specifically, all Nav channels would recover more rapidly from inactivation making more channels quickly available to participate in subsequent APs as suggested in the calculations for Figure 10D-E. A complication to this simple interpretation of the role of the slow recovery process is that any changes in the magnitude and time course of Nav currents during an AP inevitably shift the balance of Cav and Kv participation during such APs (Lingle et al., 2018), which in turn may oppose the consequences of any changes in Nav availability. Once the molecular components that participate in fast inactivation in CCs are determined, future work may permit explicit tests of the role of dual-pathway fast inactivation to regulation of firing.

### Comparing dual-pathway fast inactivation in CCs to inactivation recovery in other cells

Qualitatively and, to some extent, quantitatively similar dual-pathway fast inactivation have been described in cerebellar granule cells (Goldfarb et al., 2007), hippocampal pyramidal neurons (Venkatesan et al., 2014), and raphe neurons (Milescu et al., 2010; Navarro et al., 2020). In all cases, accumulation of Nav channels in slow recovery pathways appears to influence Nav availability and firing frequency. Although the work on granule cells and pyramidal neurons clearly demonstrate a potential role of slow recovery from fast inactivation in defining Nav channel availability and firing, the measurements do not allow assessments of the relative entry into fast and slow recovery paths from single APs.

In contrast, the work on raphe neurons exhibits both similarities and differences with the phenomenology described here. Whereas for CCs a single 5 ms depolarization drives channels into about half fast recovery and half slow recovery pathways, in the raphe neurons, only after 20% of activated channels enter slow recovery pathways and 80% fast recovery pathways (Milescu et al., 2010; Navarro et al., 2020). Experimentally, 20 Hz trains can drive raphe Nav channels into about 50% slow recovery and theoretically higher (Navarro et al., 2020), whereas in CCs Nav channels appear to be limited to about 75% occupancy in the slow recovery pathways. Although such differences might be discounted as simply being the likely consequence of different molecular underpinnings, the differences may be physiologically instructive and point to important future topics for investigation. The distribution of channels between slow and fast recovery pathways following an AP or brief depolarization might be influenced by two primary factors. One possibility is that there may be differences in the relative expression of a regulatory subunit that contributes to the slow recovery process, such that on average not all Nav channels in a cell may contain the necessary regulatory subunit. For both the raphe cells and the CCs, we think this is unlikely. In the raphe cells, modelling is consistent with the idea that channels can be driven almost completely into slow recovery pathways. For the CCs, although we were unable to drive channels into more than about 75% slow recovery, examination of the impact of fast and slow recovery rates on the state occupancies suggest that the kinetics of the recovery processes relative to the recovery intervals in the higher frequency trains can limit slow recovery pathway occupancy. A second possibility is that, although both fast inactivation pathways are entered during brief depolarization, it may be that in different types of cells, the relative rate of onset of the two types of fast inactivation processes differ. For CC, we have no direct measurements of the rates of entry into traditional fast inactivation, but that we observed 50% fast inactivated and 50% slow inactivated suggests the rates are comparable. For the raphe neurons, it is possible that the rate of normal fast inactivation is simply 4-fold faster than the inactivation leading to slow recovery. One difference in the kinetic properties of the dual-pathway processes between CCs and raphe neurons is that the fast and slow recovery time constants differ by about 20-fold in CCs (Fig. 3C), whereas in raphe neurons they differ by about 100-fold (Fig. 4A (Milescu et al., 2010)). Such a difference, whatever its molecular basis, could play a critical role in defining the dynamic range of Nav availability and firing properties in a given neuron.

### The identity of Nav currents in rat CCs

The identity of the specific Nav subtype that underlies the Nav currents in rat CCs remains unresolved. The functional data are most consistent with the idea that a single category of Nav current is found in rat CCs and this issue is addressed more directly in an associated paper (Martinez-Espinosa et al., 2020). Here we simply mention that the associated paper describes the functional similarities between rat and mouse Nav current and KO of Nav1.3 is shown to completely abolish mouse CC Nav current.

### The mechanistic basis of dual-pathway, fast inactivation in CCs

The present results have described the properties of entry into and recovery from inactivation over a variety of voltages and stimulation conditions, in order to tease apart two distinct fast inactivation processes. All manipulations point to the idea that two primary fast inactivation processes contribute to inactivation of rat CC Nav current, although the present results provide no information on the molecular components that underlie these processes. Presumably, one corresponds to traditional fast Nav inactivation, while the other remains undefined. Although the data accumulated here places constraints on the exact models that can be used to describe this behavior, quantitative description of rodent CC inactivation must await determination of the molecular determinants that underlie the two inactivation components. It should be noted that some of our observations are not fully explained by the strictly separable dual pathway model. First, we observed a slowing of fast inactivation with frequency of trains or numbers of pulses in a train. Second, although both fast and slow recovery paths are entered equivalently at strong depolarizations, near resting potentials entry and exit into those pathways are readily separable, even though none of our data support. Third, it isn’t readily apparent how relative occupancy in fast and slow pathways could be so similar over such a range of voltages, whether measured for entry or exit. Some of these issues may be related to properties of closed-state inactivation for the two pathways, for which we have no information. Furthermore, although various protocols all point to two inactivation process, it must be kept in mind that the measured time constants for these processes, although measured as two distinct exponential processes over a variety of conditions, may represent complex percolation of channels through sets of states. We anticipate that the results presented here, once the molecular determinants of the two distinct fast inactivation processes are clarified, will serve as a foundation for quantitative evaluation of the dual-pathway inactivation behavior and how it may impact on excitability.

## Acknowledgement

We thank M. Prakriya for valuable contributions during the early stages of this work. This work was supported by grant NS100695 from the National Institute of Neurological Diseases and Stroke.

---

**Fig. S1.**
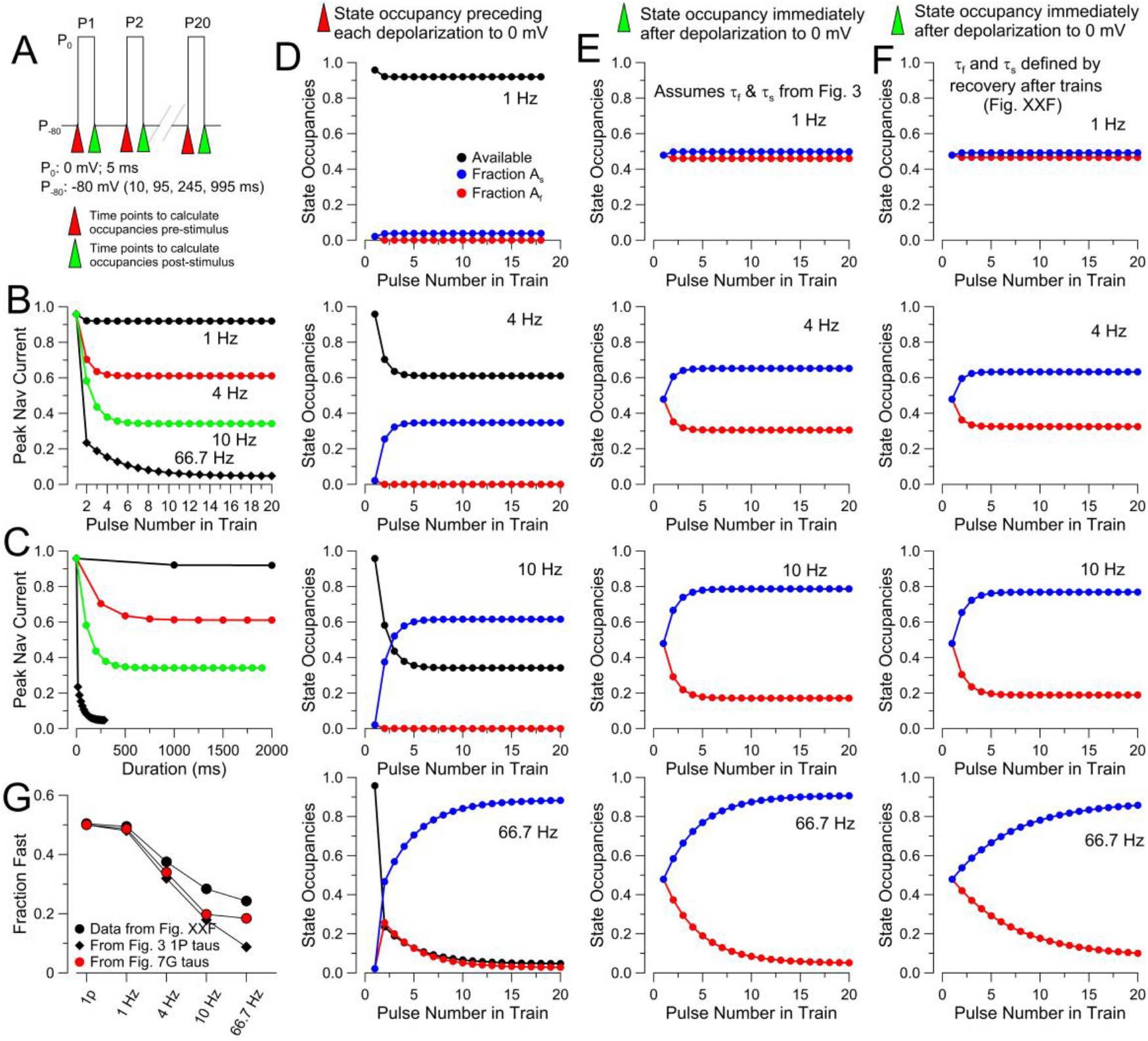
Predicted state occupancies and peak current decrement during trains of different frequencies. (A) Schematic of pulse train design and time points used for calculation of occupancies. Trains of twenty 5 ms depolarizations to 0 mV were evoked from a holding potential of −80 mV (e.g., Fig. 8), with interpulse intervals of either 10 ms (66.7 Hz), 95 ms (10 Hz), 245 ms (4 Hz), or 995 ms (1 Hz). Red arrows mark time points immediately prior to depolarizations to 0 mV at which occupancies pre-stimulus were determined. The fraction of non-inactivated channels at this point defines the effective peak current activated during the subsequent depolarization. Green arrows mark the time point immediately after termination of the depolarizing pulse. The fraction of channels in slow and fast recovering inactivated states defines the fraction of fast and slow recover components that one would predict during recovery intervals begun at these time points. To determine fractional occupancies the following assumptions and calculations were made. First, beginning from a holding potential of −80 mV and from the steady-state inactivation curve of Fig. 1E, the fraction of available channels is 0.95, with 0.025 each in slow and fast inactivated states. Second, we assume that whatever the fraction of channels activated by a depolarization, half will inactivate into fast recovery pathways and half into slow recovery pathways. Third, the fraction of channels that inactivated into each path during the 5 ms depolarization are incremented by the fraction of channels that remained in fast and slow recovery paths prior to the depolarization. These sums then define the fractions of channels in fast and slow recovery paths, if recovery was then allowed to proceed. Fourth, for each recovery interval (whether 10, 95, 245, or 995 ms), the time constants for fast and slow recovery from Fig. 3C at −80 mV were used to calculate the fraction of fast or slow inactivated channels that would be expected to recover from inactivation during that interval. This then allows determination of the fractional occupancies prior to each subsequent depolarization. (B) The decrement in peak Nav current based on determination of fractional availability prior to each 5 ms depolarization was determinated and plotted as a function of number of the pulse in a train for each of the indicated frequencies based on inactivation time constants from Fig. 3. Compare to measured decrements in depolarization-evoked Nav currents during trains in Fig. 8. (C) Replot of the values in panel B as a function of time. (D) The calculated state occupancies for channels available for activation (immediately before each 5 ms depolarization), channels in fast recovery inactvated states, and channels in slow recovery inactivated states are plotted for, from top to bottom, trains of 1, 4, 10, and 66.7 Hz. Note that for train frequencies or 1, 4, and 10 Hz, essentially all channels that inactivate into fast recovery inactivated states recover from inactivation between each depolarization, while at 66.7 Hz, there is an initial increase in the fraction of channels in fast recovery states for early pulses, which then decreases as occupancy of channels in slow recovery states increases. (E) State occupancies for channels in fast recovery pathways and slow recovery pathways immediately following the 5 ms depolarization are plotted for, from top to bottom, trains at 1, 4, 10, and 66.7 Hz, respectively. In all cases, time constants were taken from Fig. 3C (*τ_f_*=16 ms; τ_s_ = 388 ms) and used for all trains. (F) As in E, but values for *τ_f_* and τ_s_ were assumed to vary with train frequency as in Fig. 8G, for which *τ_f_* becomes slower as train frequency is increased. For train frequencies of 1, 4 and 10 Hz, the recovery intervals are sufficiently long that fractional occupancies are similar for the small differences in time constants used. At 66.7 Hz, changes in the fast recovery time constant results in substantial changes in fractional recovery during the 10 ms recovery interval, thereby slowing the decrease in occupancy of channels in fast recovery states. (G) Changes in the fraction of fast recovery following a 10 pulse train at various frequencies are plotted, showing the data plotted from Fig. 8F, along with the calculated occupancy of fast recovery states from panel E, and also the occupancies from panel F. Although peak current amplitudes during a train at 10 Hz fall to levels below 0.05, depending on apparent time constants of fast inactivation fractional occupancy of channels in fast recovery states can still be as much as 0.2, generally consistent with experimental observations (Fig. 7E, 8G).

**Fig. S2.**
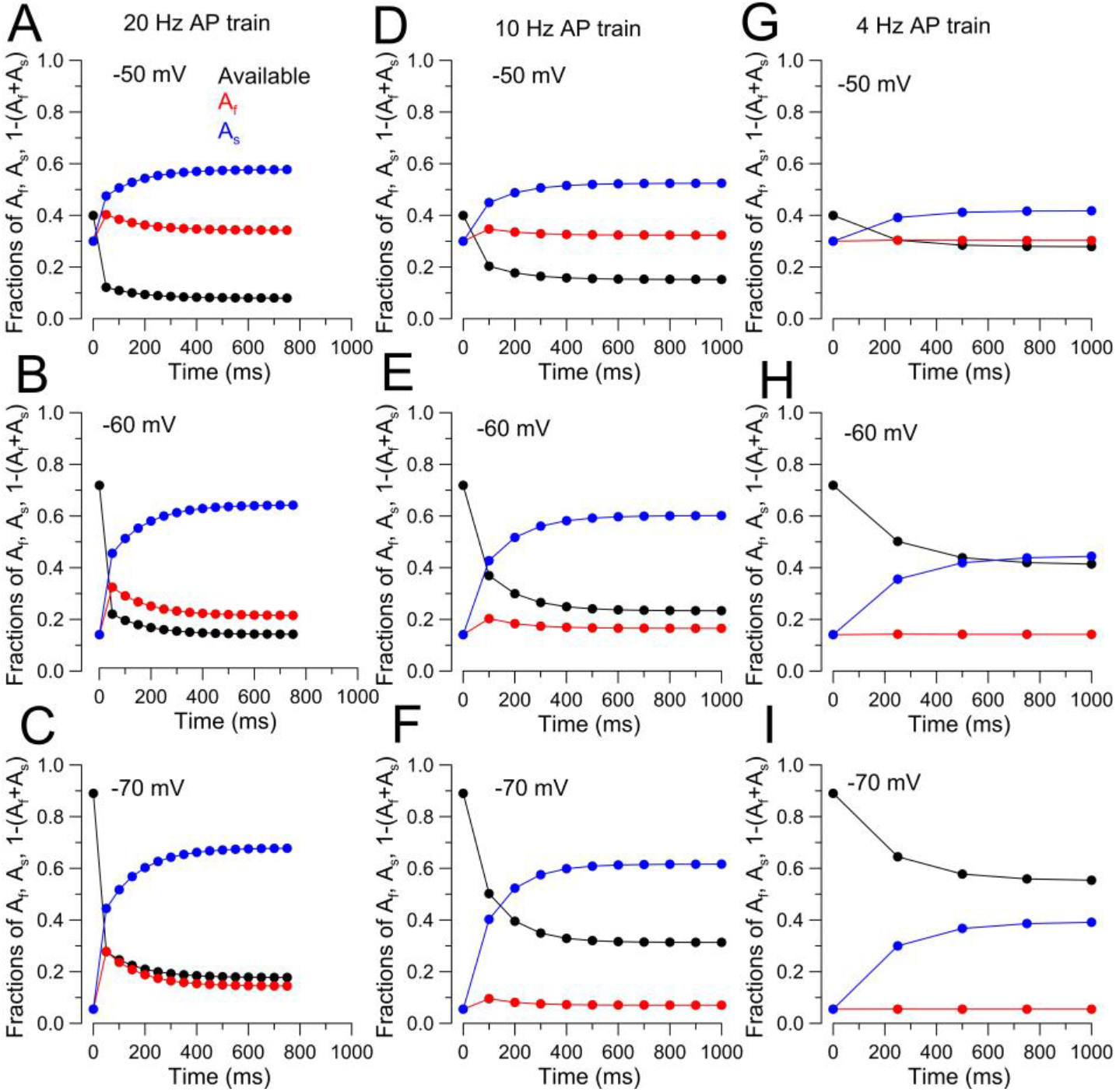
Calculated fractional occupancies of fast and slow recovery pathways during AP clamp waveform trains. Strategy for calculation of occupancies follows that in Fig. S1, with identical assumptions with initial state occupancies defined by the steady state availability curve (Fig. 1E, 0.95787 (−80), 0.890 (−70 mV), −0.719 (−60 mV), 0.400 (−50 mV)), with assumption that inactivated fraction begins evenly distributed between fast and slow recovery paths. For the impact of the AHP, we assumed a 34.6 ms AHP centered about −50 mV. For 4 Hz stimulation, there is then a 200.2 ms interval at the holding potential, for 10 Hz, there is 49.8 ms, and for 20 Hz, only 1.4 ms. Thus, for 20 Hz stimulation the recovery is dominated by the tail current period at −50 mV, rather than the holding potential. For the AP clamp waveform, there is also 5 ms at − 50 mV that precedes the upswing of the AP voltage waveform. Given the slow rates on onset or recovery from inactivation, this 5 ms interval has negligible impact on changes in occupancy of inactivated states. This procedure essentially defines the expected state occupancies immediately preceding each AP. (A-C) Calculated state occupancies for a 20 Hz AP train, for −50, −60, and − 70 mV holding potentials are shown. Following the initial AP waveform, at the time of the second AP waveform there is increased occupancy of both fast and slow recovering pathways, but with subsequent APs the fraction of channels in slow recovery pathways increases, while those in fast recovery pathways decreases. (D-F) State occupancies are plotted for the 10 Hz cases, qualitatively exhibited behavior similar to that for 20 Hz. (G-I). State occupancies plotted for the 4 Hz trains.

## Notes

### Competing Interest Statement

The authors have declared no competing interest.

